# The Detailed Organization of the Human Cerebellum Estimated by Intrinsic Functional Connectivity Within the Individual

**DOI:** 10.1101/2020.09.15.297911

**Authors:** Aihuiping Xue, Ru Kong, Qing Yang, Mark C. Eldaief, Peter Angeli, Lauren M. DiNicola, Rodrigo M. Braga, Randy L. Buckner, B.T. Thomas Yeo

## Abstract

Distinct regions of the cerebellum connect to separate regions of the cerebral cortex forming a complex topography. While key properties of cerebellar organization have been revealed in group-averaged data, in-depth study of individuals provides an opportunity to discover functional-anatomical features that emerge at a higher spatial resolution. Here functional connectivity MRI was used to examine the cerebellum of two intensively-sampled individuals (each scanned across 31 MRI sessions). Connectivity to somatomotor cortex showed the expected crossed laterality and inversion of the body maps between the anterior and posterior lobes. A surprising discovery was connectivity to the primary visual cortex along the vermis with evidence for representation of the central field. Within the hemispheres, each individual displayed a hierarchical progression from the inverted anterior lobe somatomotor map through to higher-order association zones. The hierarchy ended near Crus I/II and then progressed in reverse order through to the upright somatomotor map in the posterior lobe. Evidence for a third set of networks was found in the most posterior extent of the cerebellum. Detailed analysis of the higher-order association networks around the Crus I/II apex revealed robust representations of two distinct networks linked to the default network, multiple networks linked to cognitive control, as well as a separate representation of a language network. While idiosyncratic spatial details emerged between subjects, each of these networks could be detected in both individuals, and small seed regions placed within the cerebellum recapitulated the full extent of the spatially-specific cerebral networks. The observation of multiple networks in juxtaposed regions at the Crus I/II apex confirms the importance of this zone to higher-order cognitive function and reveals new organizational details.

## Introduction

Within its finely organized folia the human cerebellum possesses a surface area that is 4/5^th^ the size of the cerebral cortex (Sereno et al. 2020). Regions of the cerebral cortex project to specific zones of the cerebellum that, in turn, project back to the same regions of the cortex (Schmahmann and Pandya 1997; Strick, Dum and Fiez 2009). While historical focus has been on motor function, the cerebellum is now known to contain major portions linked to cognitive and affective functions (Leiner et al. 1986; 1989; 1993; Schmahmann and Sherman 1998; Strick, Dum and Fiez 2009; Buckner 2013; Keren-Happuch et al. 2014; Schmahmann et al. 2019; Diedrichsen et al. 2019). Over the past decade, the field has moved forward to explore the organization of cerebellar association zones with an increasing level of detail (Stoodley and Schmahmann 2009; Buckner et al. 2011; Guell et al. 2018a; King et al. 2019), including novel tools for visualization on the surface (Diedrichsen and Zotow 2015; Guell et al. 2019). What is converged upon is that between the anterior and posterior somatomotor representations is a large region linked to multiple cerebral association networks. The present work builds on these foundations by exploring the detailed organization of the human cerebellum within intensively sampled individuals. A major challenge arising in the study of cerebrocerebellar organization is that there are no direct monosynaptic connections between the cerebellum and the cerebral cortex. The afferent inputs connect via the pons and the efferents project through the deep cerebellar nuclei and thalamus (Schmahmann and Pandya 1997; Strick, Dum and Fiez 2009). This creates a formidable barrier: cerebrocerebellar circuits cannot be mapped by the mainstay historical technique of systems neuroscience – monosynaptic retrograde and anterograde tract tracing. Polysynaptic tracing techniques allowed for a breakthrough which revealed a closed-loop architecture between the cerebrum and cerebellum that included parallel motor and prominent prefrontal connections in the macaque (e.g., Middleton and Strick 2001).

Human functional connectivity MRI based on spontaneous correlation provides another critical approach to map cerebrocerebellar organization (Biswal et al. 1995; Fox and Raichle 2007). For its many weaknesses and caveats (Buckner et al. 2013; Murphy et al. 2013; Smith et al. 2013; Power et al. 2014), functional connectivity MRI is powerful for mapping cerebellar organization because it is sensitive to polysynaptic circuitry and also because it allows for a wide field of view that can cover the entire brain. Numerous cerebral locations can be simultaneously mapped to numerous cerebellar locations. Use of functional connectivity MRI for cerebrocerebellar mapping has been supported by demonstrating (1) sensitivity, in detecting the well-established (Adrian 1943; Snider and Stowell 1944) anterior and posterior somatomotor representations (Habas et al. 2009; Krienen and Buckner 2009); (2) specificity, by mapping the orderly topography of the inverted and upright body maps (Buckner et al. 2011); (3) sufficiency, by establishing cerebral stimulation can evoke a polysynaptic cerebellar response in non-human primates (Matsui et al. 2012); and (4) dependency, on intact anatomical circuits via analysis of hemipons lesions that disrupt the crossing cerebrocerebellar projections (Lu et al. 2011). Functional connectivity MRI is the basis of the present investigations.

A second challenge arises because of the small size of the human cerebellum and the limited sensitivity and resolution of human neuroimaging approaches. Most prior studies of the human cerebellum relied on group-averaging of data to boost signal-to-noise at the expense of resolution (e.g., Stoodley and Schmahmann 2009; Buckner et al. 2011; Guell et al. 2018a; King et al. 2019). What has emerged is a rough topographic organization of the cerebellum including multiple representations of sensory-attention networks and higher-order networks between the anterior and posterior lobe representations of the somatomotor map. The region surrounding the Crus I/II border is particularly intriguing because it contains representations of multiple distinct association networks. However, the detailed organization of the cerebellum, including mapping in relation to recently resolved higher-order association networks in the cerebral cortex, has been limited because spatial features are blurred together in group-averaged data (e.g., Fedorenko et al. 2012; Laumann et al. 2015; Huth et al. 2016; Braga and Buckner 2017; Gordon et al. 2017; Braga et al. 2019; Gordon et al. 2020).

Of particular relevance to the present investigations are the higher-order association networks that have been mapped within individuals in great detail for the cerebral cortex. Multiple parallel networks have been discovered and replicated that possess neighboring regions that are often closely juxtaposed on the same sulcus (e.g., Braga et al. 2019). These multiple distinct juxtaposed networks are hypothesized to support functions related to remembering, social inferences, and language – some of the most advanced forms of human thought (e.g., Fedorenko et al. 2012; Braga and Buckner 2017; Buckner and DiNicola 2019; DiNicola et al. 2020; Gordon et al. 2020; Braga et al. 2020). The present work focuses on these higher-order association networks within the individual by first identifying them in the cerebral cortex and then using the detailed characterization to map the cerebellum.

Providing a foundation, Marek et al. (2018) recently mapped the functional organization of the cerebellum from an openly available data resource of 10 highly sampled individuals (Gordon et al. 2017). This work illustrates the opportunity and challenges of precision cerebellar mapping. First, they found considerable variability between individuals that likely represents a combination of neuroanatomical variance as well as technical variance attributed to mapping at the edge of resolution and sensitivity limits. Second, they discovered organizational details that would be difficult to appreciate in group-averaged data. For example, they identified large representations of cognitive-control networks as well as attentional networks that have been underemphasized in prior group-averaged analyses. Finally, their individualized analyses confirmed the complexity of the cerebellar apex near the Crux I/II border with multiple association networks converging near to one another in most individuals.

Here we extend precision cerebellar mapping to investigate the detailed functional organization of the cerebellum in two intensively sampled individuals. Particular focus is placed on the higher-order association networks that have been recently disentangled in the cerebral cortex including multiple parallel networks linked to the default network (Braga et al. 2017; 2019; DiNicola et al. 2020) and also their separation from a juxtaposed left-lateralized language network (Braga et al. 2020). While not an original intent of our explorations, the high-resolution mapping within the individual yielded evidence for a visual representation near the vermis as predicted by van Es and colleagues (2019).

## Methods

### Overview

The present explorations characterized the functional organization and topography of the cerebellum of two intensively-studied individual subjects using data from 31 resting-state functional MRI (fMRI) sessions. Many of the analyses recapitulated strategies used previously to examine cerebellar organization in group-averaged data (Buckner et al. 2011), applied here to the individual. First, we examined the cerebrocerebellar functional connectivity of the somatomotor cortex to detect the well-established contralateral connectivity (establishing sensitivity) and topographic organization of the body representations (establishing specificity). Next, we estimated individual-specific cerebral parcellations using a Bayesian hierarchical approach (Kong et al. 2019) with data from the cerebellum blinded. We then used the cerebral network estimates as the anchor to comprehensively map the topographic organization of the cerebellum. All critical analyses were replicated between independent datasets within the same individual and generalized between the two individuals. Having established estimates of within-individual cerebellar organization using a cerebral-driven approach, we reversed the analysis strategy to validate the findings using a cerebellar-driven approach. Small seed regions were defined within the cerebellum and the functional connectivity patterns were visualized in the cerebral cortex to determine whether the cerebellar regions were coupled to the cerebral cortex with spatially-specific patterns. This same approach was also used to determine whether the multiple cerebellar representations would display the same patterns. As a final analysis, we explored in detail a surprising observation surrounding the identification of a zone of the cerebellar vermis that was linked to early retinotopic visual cortex.

### Participants

The two subjects in this study were healthy right-handed adult women (age 22 and 23) recruited from the greater Boston community and screened to exclude a history of neurological and psychiatric illness or ongoing use of psychoactive medications. The data from this study has been previously utilized in another study that focused on the cerebral cortex (Braga et al. 2019).

### MRI data acquisition

The resting-state fMRI data were collected at the Harvard Center for Brain Science on a Siemens Prisma-Fit 3T MRI scanner using the vendor’s 64-channel phased-array head-neck coil (Siemens, Erlangen, Germany). For both participants, 31 MRI sessions were collected with at least 2 resting-state blood oxygenation level-dependent (BOLD) runs included in each session. In total, 63 runs were obtained for each participant within 40 weeks. Each run lasted 7 min and 2 sec. Foam padding on the top and side of the head helped immobilize the head and reduce motion. Participants were instructed to remain still, stay awake, and maintain fixation on a centrally presented black crosshair with a light grey background viewed through a mirror. The scanner room was illuminated to enhance alertness.

BOLD fMRI (Kwong et al. 1992; Ogawa et al. 1992) data were acquired using a multiband gradient-echo echo-planar pulse sequence (Setsompop et al. 2012): TR = 1000 ms, TE = 32.6 ms, flip-angle = 64°, 2.4 mm isotropic voxels, matrix 88 × 88 × 65, multislice 5×acceleration, 422 frames for each run (with 4 frames then removed for T1 equilibration). The custom sequence was generously provided by the Center for Magnetic Resonance Research (CMRR) at the University of Minnesota as implemented for the Human Connectome Project (HCP; Xu et al. 2012; Van Essen et al. 2013). Signal dropout was minimized by automatically selecting a slice plane 25° from the anterior-posterior commissural plane toward the coronal plane (van der Kouwe et al. 2005; Weiskopf et al. 2006; Mennes et al. 2014). A T1-weighted structural image was obtained in each session using a rapid multiecho magnetization-prepared rapid gradient echo (MPRAGE) three-dimensional sequence (van der Kouwe et al. 2008): TR = 2200 ms, TE = 1.57, 3.39, 5.21, 7.03 ms, TI = 1100ms, flip-angle = 7°, voxel size 1.2 mm, matrix 192 × 192 × 176, in-plane generalized auto-calibrating partial parallel acquisition (GRAPPA) acceleration 4 (see Nielsen et al. 2019 for analysis of this rapid variant). A dual gradient-echo B0 field map was acquired to correct for susceptibility-induced gradient inhomogeneities: TE = 4.45, 6.91 ms with slice prescription/spatial resolution matched to the BOLD sequence (Braga et al. 2019).

There were two exclusion rules for quality control of the BOLD data: (1) maximum absolute motion more than 2 mm and (2) slice-based temporal signal-to-noise ratio ≤130. Following these rules, 1 run was excluded for Subject 1, and 2 runs were excluded for Subject 2. Overall, 62 runs were usable for Subject 1 and 61 runs for Subject 2 (thereby recovering some data that failed processing in Braga et al. 2019).

### Data processing

Data were processed through an in-house preprocessing pipeline (“iProc”; Braga et al. 2019) that combined tools from FreeSurfer (Fischl 2012), FSL (Jenkinson et al. 2012) and AFNI (Cox et al. 1996; 2012). A key goal of this processing pipeline is to align data within an individual across runs and sessions with a single interpolation (to minimize blurring) and to high-resolution final targets (1mm). The two participants were processed independently. For each participant, a mean BOLD template was created by taking the mean of all fieldmap-unwarped middle volumes after registration to an upsampled (1.2-mm), unwarped mid-volume template. Five different transformation matrices were calculated to align each BOLD image volume: (1) a motion correction matrix for each volume to the run’s middle volume, (2) a matrix for fieldmap-unwarping the run’s middle volume, (3) a matrix for registering the fieldmap-unwarped middle volume to the mean BOLD template, (4) a matrix for registering the mean BOLD template to the subject’s 1-mm native space T1 image, and (5) a matrix for registering the native space T1 to the MNI152 1-mm atlas. The first four transformation matrices were composed into one single matrix, and all five transformation matrices into another. The composed matrices were used to project, in parallel, each original BOLD volume to the native space T1 or to MNI space, each in a single interpolation to reduce spatial blur (Braga and Buckner 2017; Braga et al. 2019).

Motion nuisance parameters, whole brain, ventricular, and deep white matter signal were calculated and regressed out from the native space and MNI-space-projected data. A 0.01-0.1 Hz bandpass filter was applied to the data. In the case of the cerebral cortex, the native space data were projected to the fsaverage6 standardized cortical surface mesh with 40,962 vertices per hemisphere (Fischl et al. 1999). A 2mm full width at half-maximum (FWHM) kernel was used for surface smoothing. In the case of the cerebellum, the volumetric data in MNI space were smoothed with a 4mm FWHM kernel. A manually corrected mask was used to extract the cerebellum (Fig. 1).

**Fig. 1.**
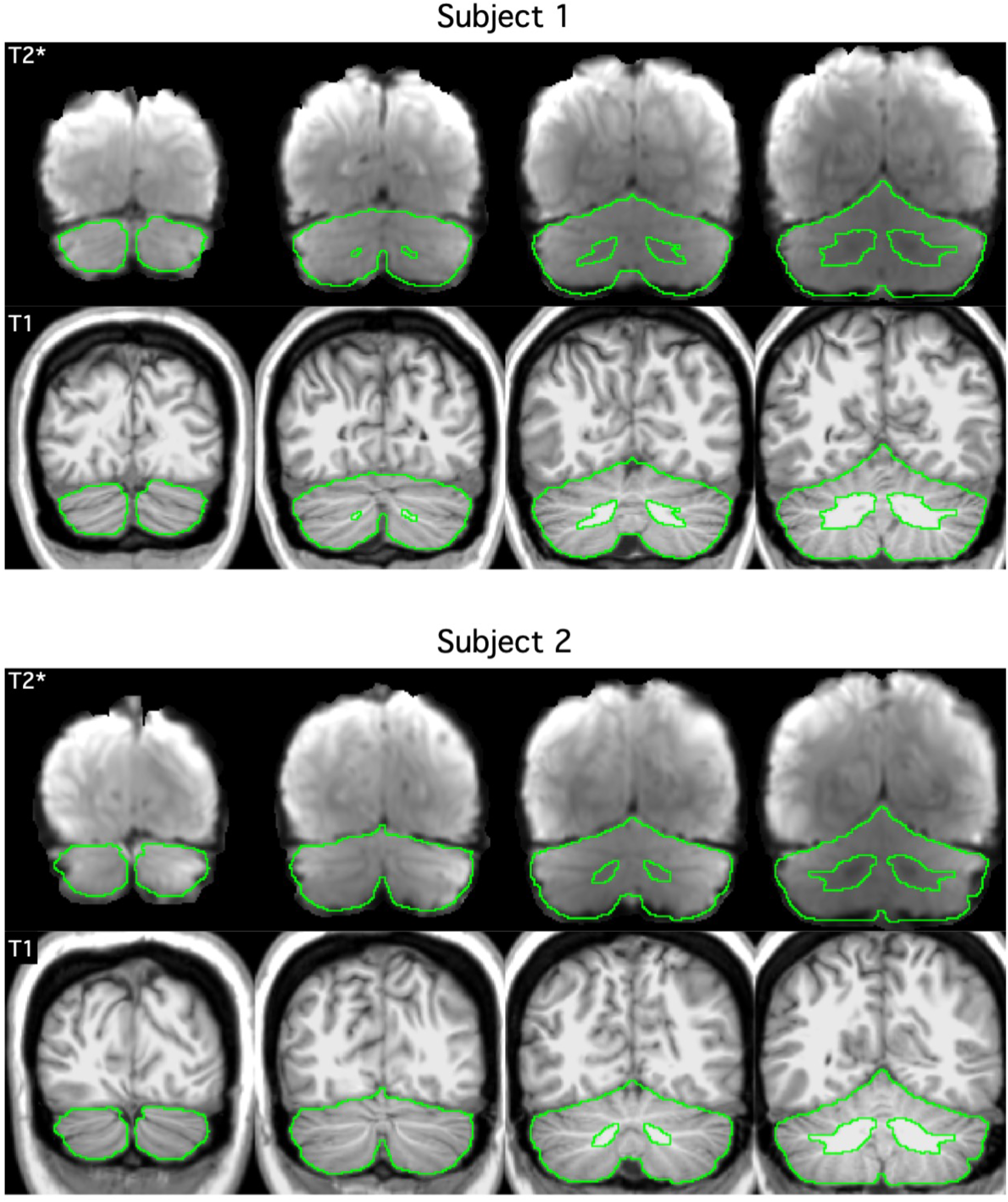
Extracted cerebellar boundaries for Subjects 1 and 2 in MNI152 space. The top row of each panel displays the T2* BOLD images. The bottom row of each panel shows the T1-weighted images. The green line shows the boundary of the cerebellum mask manually delineated based on each individual subject’s T1-weighted image. There is good agreement between the green line and the cerebellum in the T2* images, suggesting good T1-T2* alignment. Imperfections in the BOLD data are also visible, e.g., signal dropout near the green cerebellar boundary.

### Visualization on the cerebellar surface

For many analyses the cerebral data are visualized on the cerebral surface and the cerebellar data within the cerebellar volume. In addition to volume representation within the cerebellum, we also adopted a modeled projection of the data onto an estimated cerebellar surface. More specifically, the volumetric parcellations were projected to a cerebellar flatmap using the spatially unbiased infratentorial template (SUIT) toolbox (http://www.diedrichsenlab.org/imaging/suit_fMRI.htm). An outer (grey-matter) cerebellar surface and inner (white-matter) cerebellar surface were defined by Diedrichsen and Zotow (2015). Each vertex on the flatmap has a corresponding vertex on the outer surface and a corresponding vertex on the inner surface. To project the parcellations from the volume to the flatmap, each vertex on the flatmap was assigned to the most frequent network of the voxels along the line connecting the corresponding vertices on the outer and inner surfaces. Given the current resolution, the cerebellar surfaces are unable to reflect the highly fine cerebellar folia. Therefore, the flatmap is not a true unfolding of the cerebellar cortex. Nevertheless, based on our manual inspection, the flatmap was able to capture the main parcellation features.

### Discovery and replication data sets

For each individual subject, the 31 sessions were divided into discovery and replication sets based on odd and even sessions. There were 16 sessions in the discovery set and 15 sessions in the replication set. For Subject 1, there were 32 runs in the discovery set and 30 runs in the replication set. For Subject 2, there were 32 runs in the discovery set and 29 runs in the replication set.

### Motor hand region functional connectivity

To establish that our analysis procedures are *sensitive* to detect known organizational features of the cerebellum, we computed functional connectivity between cerebral motor regions positioned within or near the hand representation and the cerebellum. We estimated the motor hand region seed region location from Buckner et al. (2011) defined in MNI152 space. More specifically, we considered 6mm-radius seed regions centered at MNI152 coordinates (x = −41, y = −20, z = 62) for the left-hand region and (41, −20, 62) for the right-hand region. For each run of each subject, we computed the Pearson’s correlation (i.e., functional connectivity) between the time courses of the seed regions and all cerebellar voxels. The functional connectivity values were averaged across all runs after Fisher’s r-to-z transformation. We then computed the difference between the functional connectivity maps of the left-hand and right-hand seed regions for each individual (paralleling the strategy established in Krienen and Buckner 2009).

### Foot, hand and tongue representations

To establish *specificity* to detect spatially separate features within the cerebellum, we explored the topography of the different body parts in each individual’s cerebellum by leveraging their long-established spatial arrangement (Adrian 1943; Snider and Sowell 1944) and previous identification in group-level cerebellar analyses in people (e.g., Buckner et al. 2011; Diedrichsen and Zotow 2015). The group-level coordinates from Buckner et al. (2011) did not work well for the foot and tongue representations, potentially due to inter-individual differences not captured at the group level. Therefore, we considered a strategy of picking individual-specific seed regions. More specifically, we manually picked single vertices representing hand, foot and tongue regions on the fsaverage6 surface based on functional connectivity within the discovery set. The original MNI152 coordinates of foot, hand and tongue regions (Buckner et al. 2011) were used to guide our selection. Functional connectivity was then computed using the replication set. For each run of each subject within the replication set, we computed the Pearson’s correlation between the time series of the motor seed regions and all cerebellar voxels. We then binarized the functional connectivity maps for each body part and overlaid them to visualize the somatomotor organization. Thresholds were set at z ≥ 0.1 for foot, z ≥ 0.2 for hand, and z ≥ 0.1 for tongue.

### Individual-specific parcellation of the cerebral cortex

Having established that functional connectivity was able to detect and map the somatomotor topography within each individual’s cerebellum, we moved on to characterize the individual-specific organization of cerebrocerebellar coupling across the entire cerebral cortex. We started by estimating the individual-specific cerebral cortical parcellations of the two individuals based on the connectivity profiles of each cortical region (vertex). Following our previous study (Yeo et al. 2011), the connectivity profile of a cortical vertex was defined as its functional coupling to 1175 regions of interest (ROIs). These ROIs consist of single vertices uniformly distributed across the surface meshes. For each participant, the Pearson’s correlation was computed between the fMRI time series at each vertex (40,962 vertices per hemisphere) and the 1175 ROIs. The 40962 × 1175 correlation matrix of each hemisphere was then binarized by keeping the top 10% of the correlations to obtain the functional connectivity profiles (Yeo et al. 2011).

A multi-session hierarchical Bayesian model (MS-HBM) was then utilized for estimating cortical networks in the two individuals (Kong et al. 2019). The MS-HBM was initialized with a 10-network group-level parcellation estimated from the HCP S900 data release using the group-level clustering algorithm from Yeo and colleagues (Yeo et al. 2011; Kong et al. 2019). We note that the cerebral cortex is organized in a hierarchical fashion, so there is not a single optimal resolution in terms of the number of networks (Power et al. 2011; Yeo et al. 2011). In this study, we consider the 10-network resolution because the network topography within the cerebellum was difficult to interpret with more networks. However, the 10-network resolution was sufficiently fine to distinguish default network A from default network B, and also identify the distinct language network, which were critical targets (Braga and Buckner 2017; Kong et al. 2019; Braga et al. 2020). Thus, this choice reflected a balance to detect known target networks but also to limit the dimensionality of the solution to a tractable level.

To test the reliability of the individual-specific cerebral cortical parcellations, the MS-HBM algorithm was independently applied to the discovery and replication sets. Once reliability was established, we obtained the best estimate of the cerebral cortical parcellations for each individual, by applying the MS-HBM to all resting-state fMRI sessions (similar to Yeo et al. 2011).

### Individual-specific cerebellar parcellation

The individual-specific cerebral cortical parcellations (from the previous section) were used to estimate individual-specific cerebellar parcellations using a winner-take-all algorithm (Buckner et al. 2011; Marek et al. 2018). To test the reliability of the individual-specific cerebellar parcellations, the same procedure was independently repeated for the discovery and replication sets. Finally, to obtain the best estimate of the cerebellar parcellations for each individual, the procedure was also applied to all resting-state fMRI sessions.

Specifically, focusing on the analysis as applied to the discovery set, for each run of each subject, we computed the functional connectivity (i.e., Pearson’s correlation) between the time courses of all cerebellar voxels and all cerebral cortical vertices. The functional connectivity values were averaged across all runs for each subject within the discovery set. For each cerebellar voxel, we identified the top 400 cerebral cortical vertices with the strongest correlations with the voxel’s fMRI time course. Recall that all cortical vertices had been assigned to a cerebral cortical network by the HS-HBM analysis (see previous section). The cerebellar voxel was then assigned to the cerebral cortical network that was most frequently present among the 400 vertices. Experiments (not reported) using different number of vertices (100 to 1000 vertices) yielded highly similar cerebellar parcellations, so here we will focus on the parcellations generated with 400 vertices. The procedure was applied to the replication set independent of data from the discovery set and then, after establishing reliability, was applied to all of the data simultaneously to obtain the best estimate of the cerebellar parcellation (similar to Buckner et al. 2011).

### Cerebellar-initiated functional connectivity

The above analyses yield cerebellar parcellations that are dependent on the assumptions of the cerebral network parcellation, including the decision of how many distinct networks to estimate in the MS-HBM. To provide an alternative approach anchored directly from the cerebellum, we also explored the cerebral correlation patterns when beginning the analyses with small seed regions placed in the cerebellum. Such a strategy allows the specificity of the cerebral pattern to be observed without *a priori* assumptions about the distributed organization within the cerebrum or the number of networks in the parcellation solution. Positive evidence that the parcellations reflect stable organizational properties arises if the cerebral networks generated from small cerebellar seed regions recapitulate the estimated cerebral networks in an anatomically-specific manner. Negative evidence is obtained when the generated networks are orthogonal or non-specific. Moreover, multiple representations of the same network can be examined in the cerebellum to explore whether the multiple candidate networks zones do, in fact, each independently correlate specifically with the same cerebral network (e.g., anterior and posterior cerebellar representations of the language network).

To further remove bias from this set of confirmation analyses, seed regions were manually selected from the default, language, visual and dorsal attention networks based on functional connectivity within the discovery set. The final set of functional connectivity maps were computed using the replication set.

### Estimation of V1 from histology and functional connectivity gradients

As the results will reveal, a surprising finding was a region of the cerebellar vermis that was associated with the visual network linked to early retinotopic visual cortex. To explore the details of this finding further we estimated the location of primary visual cortex (V1) within each subject using both histology and functional connectivity gradients. The V1 histology was performed by the Juelich group (Amunts et al. 2000) and then projected to FreeSurfer fsaverage space (Fischl et al. 2008). Functional connectivity gradients were used to detect abrupt local changes in functional connectivity. Previous studies have suggested the feasibility of functional connectivity gradients to localize the V1 boundary in-vivo (Wig et al. 2014; Laumann et al. 2015; Gordon et al. 2016).

### Quantitative relationship between cerebral and cerebellar network representations

Our previous study (Buckner et al. 2011) suggested that larger cerebral cortical networks have larger representations within the cerebellum at the group-level, with some hints of disproportionate representation of certain networks (e.g., the frontoparietal control network). Marek et al. (2018) performed an extensive analysis of the relationship between cerebral and cerebellar network representations within individuals revealing evidence for disproportionate frontoparietal representation. Motivated by these prior analyses, we conducted a similar analysis here but, unlike the previous work, explicitly included a clear candidate language network as well as separately calculated the relationship for the left and right cerebellar hemispheres, given the known laterality differences for certain networks (e.g., language). Specifically, for each cerebellar hemisphere, we computed the percentage of voxels assigned to each network within the cerebellar mask in MNI space and the percentage of vertices assigned to each network on the contralateral cerebral surface. We then compared the percentage of cortical surface dedicated to each network with the percentage of cerebellar volume dedicated to the same network in the contralateral hemisphere.

## Results

### Cerebral motor regions show robust functional connectivity with the contralateral cerebellum within the individual

Subtraction of functional connectivity seed maps between the left-hand and the right-hand cerebral cortical seed regions revealed contralateral connectivity of the motor representations in the cerebellum for both individuals (Fig. 2). Both primary and secondary hand-region representations in the anterior and posterior lobes of the cerebellum were evident.

**Fig. 2.**
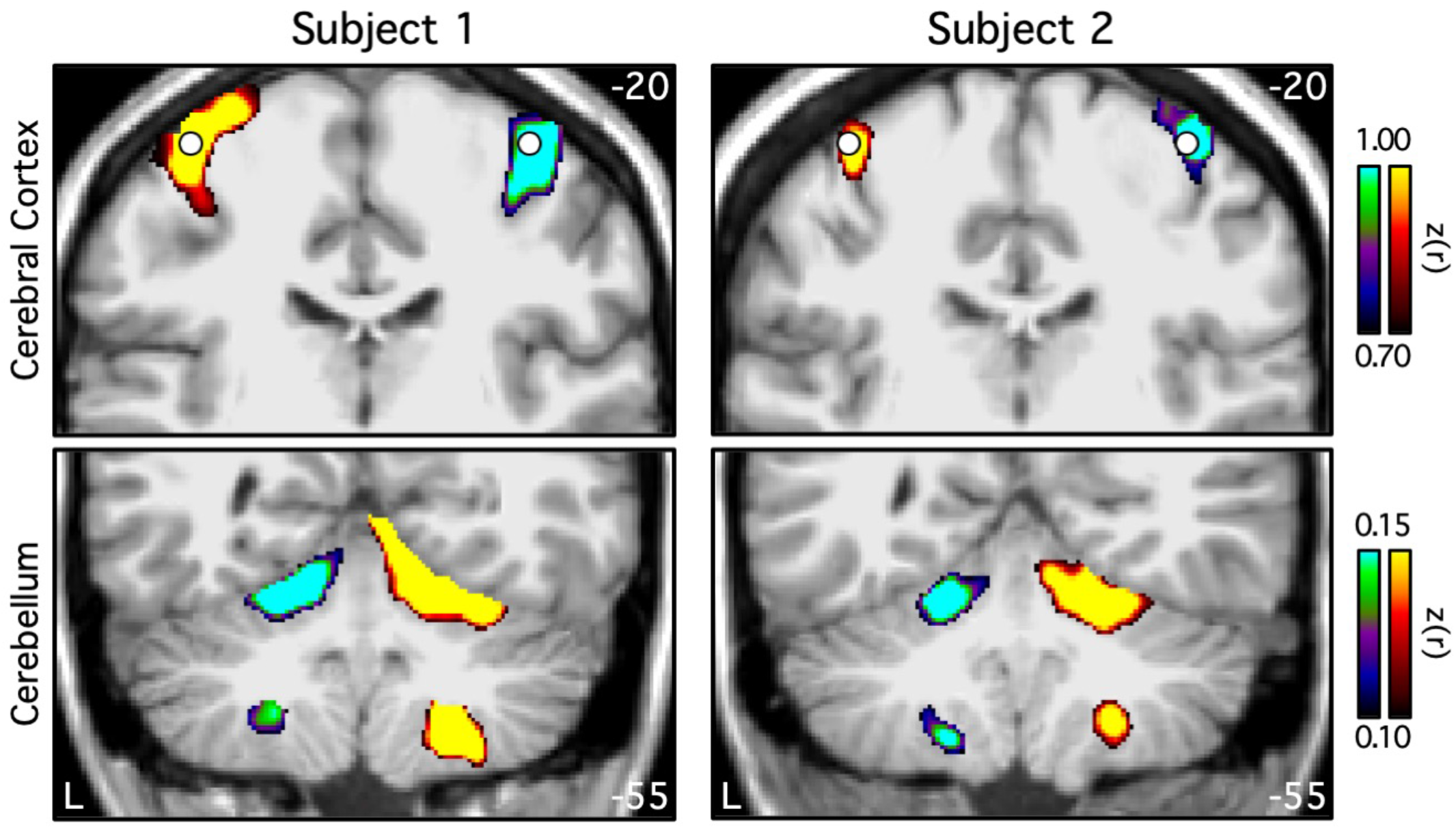
Functional connectivity of the cerebral cortical motor hand regions reveals contralateral somatomotor representations in the cerebellum of individual subjects. Coronal sections (top: y = −20; bottom: y = −55) display differences between functional connectivity of seed motor regions for each individual subject. Seed locations at or near the left and right hand motor representations are indicated by white circles. Warm colors show the connectivity of the right seed region subtracted from the connectivity of the left seed region. Cool colors represent the reverse subtraction. Color bars indicate correlation strength [z(r)]. The left hemisphere is displayed on the left. These results suggest that functional connectivity possesses *sensitivity* to detect known somatomotor representations within the individual.

### Somatomotor topography of the cerebellum is evident within the individual

Consistent with previous work on population-level cerebellar functional connectivity maps (Buckner et al. 2011; Diedrichsen and Zotow 2015), individual-specific functional connectivity maps of cerebral seed regions in estimated foot, hand and tongue representations showed an inverted topography in both individuals in the anterior lobe and a separate upright topography in the posterior lobe (Fig. 3). Although the size and location of these binarized individual representations were not exactly the same, the expected topographic ordering was consistent across both participants (see also Marek et al. 2018). A small detail was further noted. There was a spatial gap between the estimated hand and foot representations for both the anterior and posterior lobe maps. This gap, which is less evident in group-based analyses, may reflect portions of the cerebellar map associated with portions of the body between the hand and foot that are not captured fully by our limited sampling of body parts (see Saadon-Grosman et al. 2015).

**Fig. 3.**
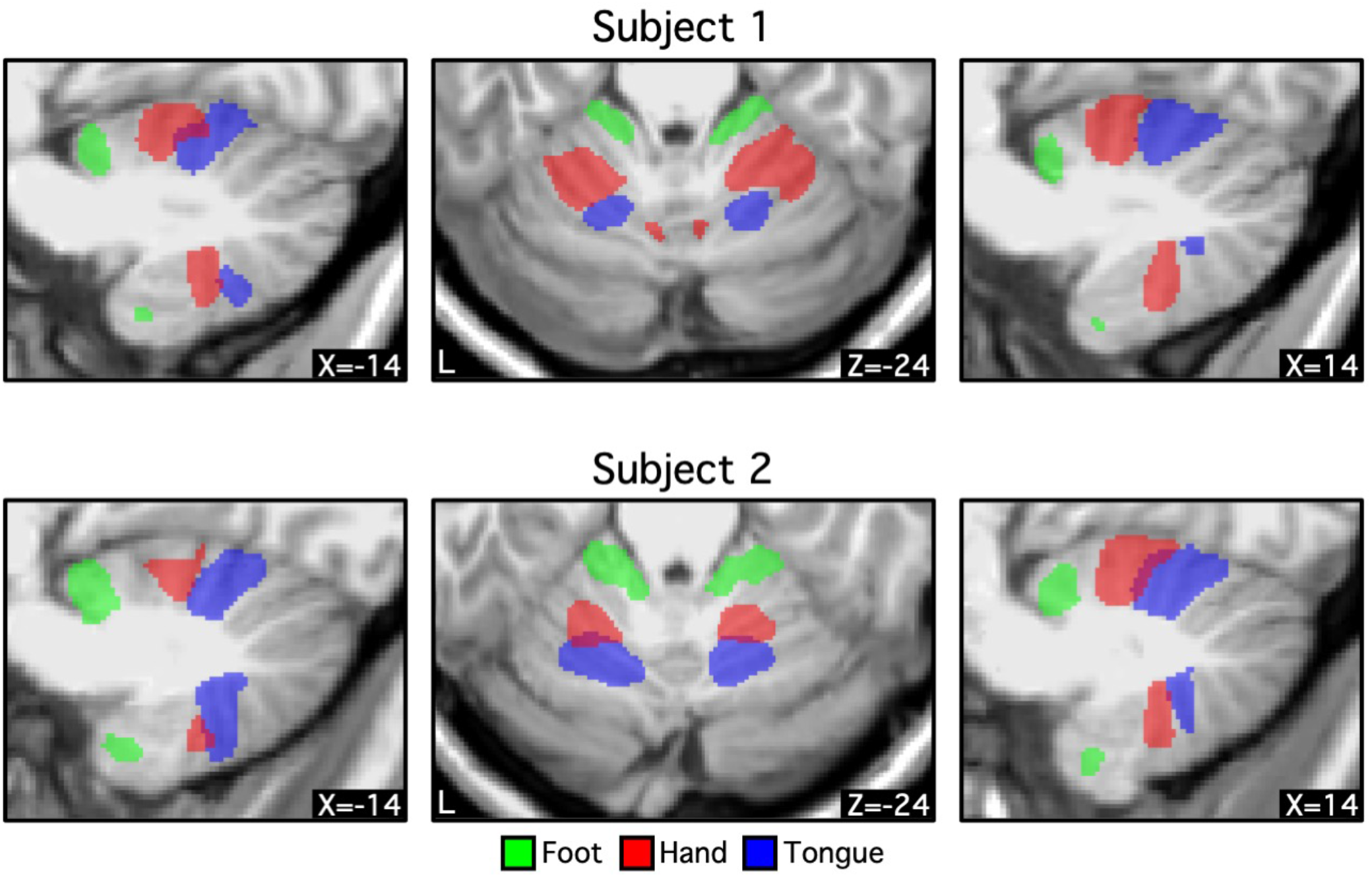
Foot-hand-tongue representation in the cerebellum revealed by functional connectivity for individual subjects. Three different bilateral cortical regions were selected on the cerebral surface based on the discovery set (Table 1). Functional connectivity of these regions was computed using the replication set. Thresholds were set at z ≥ 0.1 for foot, z ≥ 0.2 for hand, and z ≥ 0.1 for tongue. Green, red, and blue colors represent foot, hand and, tongue, respectively. Coordinates at the bottom right indicate the section level in MNI152 space. In both individuals, the order of the somatomotor representation in the anterior lobe is foot-hand-tongue, while the order in the posterior lobe is inverted. Note the consistent spatial gap between the foot and hand representations that may reflect the intervening body representation. These results suggest that functional connectivity possesses *specificity* to detect the known spatial topography of somatomotor representations within the individual.

### Cerebral cortical and cerebellar parcellations are reliable within individuals

For each subject, the 31 sessions were split into discovery and replication sets based on odd and even sessions. The MS-HBM (Kong et al. 2019) was applied to the discovery and replication sets independently to derive 10-network cerebral cortical parcellations (Fig. 4). The parcellations within each subject were highly similar across discovery and replication sets. In Subject 1, 91.4% of cortical vertices were assigned to the same networks across discovery and replication sets. In Subject 2, 90.2% of cortical vertices were assigned to the same networks across discovery and replication sets. By contrast, overlaps between the cortical parcellations of Subjects 1 and 2 were 59.8% in the discovery set and 60.4% in the replication set. Thus, between-subject variability was substantially greater than within-subject variability.

**Table 1.** Locations of seed regions used to create somatomotor topography.

|  | Subject 1 | Subject 2 |
| --- | --- | --- |
| Left hemisphere |  |  |
| Foot | 2110 | 24883 |
| Hand | 5768 | 5768 |
| Tongue | 32897 | 32897 |
| Right hemisphere |  |  |
| Foot | 34531 | 17843 |
| Hand | 6167 | 6167 |
| Tongue | 12417 | 12417 |

The number represents the index of each hemisphere on the fsaverage6 surface. Note that vertex index starts from 1 (rather than 0).

**Fig. 4.**
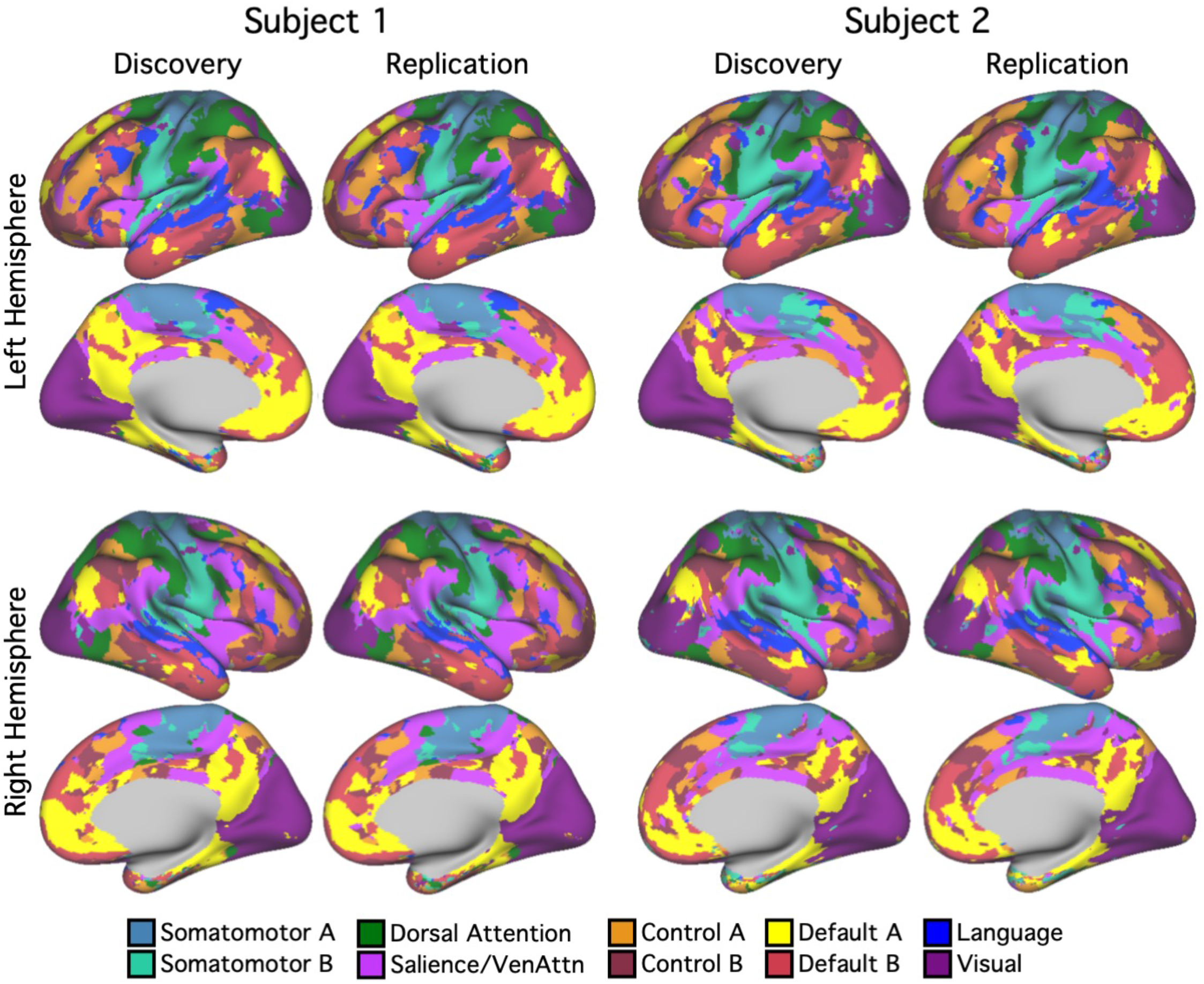
Cerebral cortical network parcellations were highly reliable across discovery and replication sets within individuals. 10-network individual-specific cerebral network parcellations were estimated by applying a multi-session hierarchical Bayesian model (Kong et al. 2019) to the discovery (16 sessions) and replication (15 sessions) datasets independently. The individual-specific cortical parcellations were replicable within subjects. In Subject 1, 91.4% cortical vertices were assigned to the same networks across discovery and replication datasets. In Subject 2, 90.2% cortical vertices were assigned to the same networks across discovery and replication datasets. Networks are colored as labelled in the bottom legend using network names common in the literature for convenience. The names should not be taken to mean that networks code solely for functions associated with their assigned names, which are often derived from an evolving literature.

For the cerebellar parcellation, we applied a winner-takes-all algorithm (Buckner et al. 2011; Marek et al. 2018) based on the functional connectivity between the cerebral cortex and the cerebellum. Cerebellar parcellations were estimated from the discovery and replication sets independently (Fig. 5). The cerebellar parcellations within each subject were highly similar across discovery and replication sets. Overlap between parcellations in the discovery and replication sets were 83.8% for Subject 1 and 84.2% for Subject 2. By contrast, overlap between the cerebellar parcellations of Subjects 1 and 2 were 42.8% in the discovery set and 42.8% in the replication set. Thus, between-subject variability was substantially greater than within-subject variability for the cerebellum as well as cerebral cortex.

**Fig. 5.**
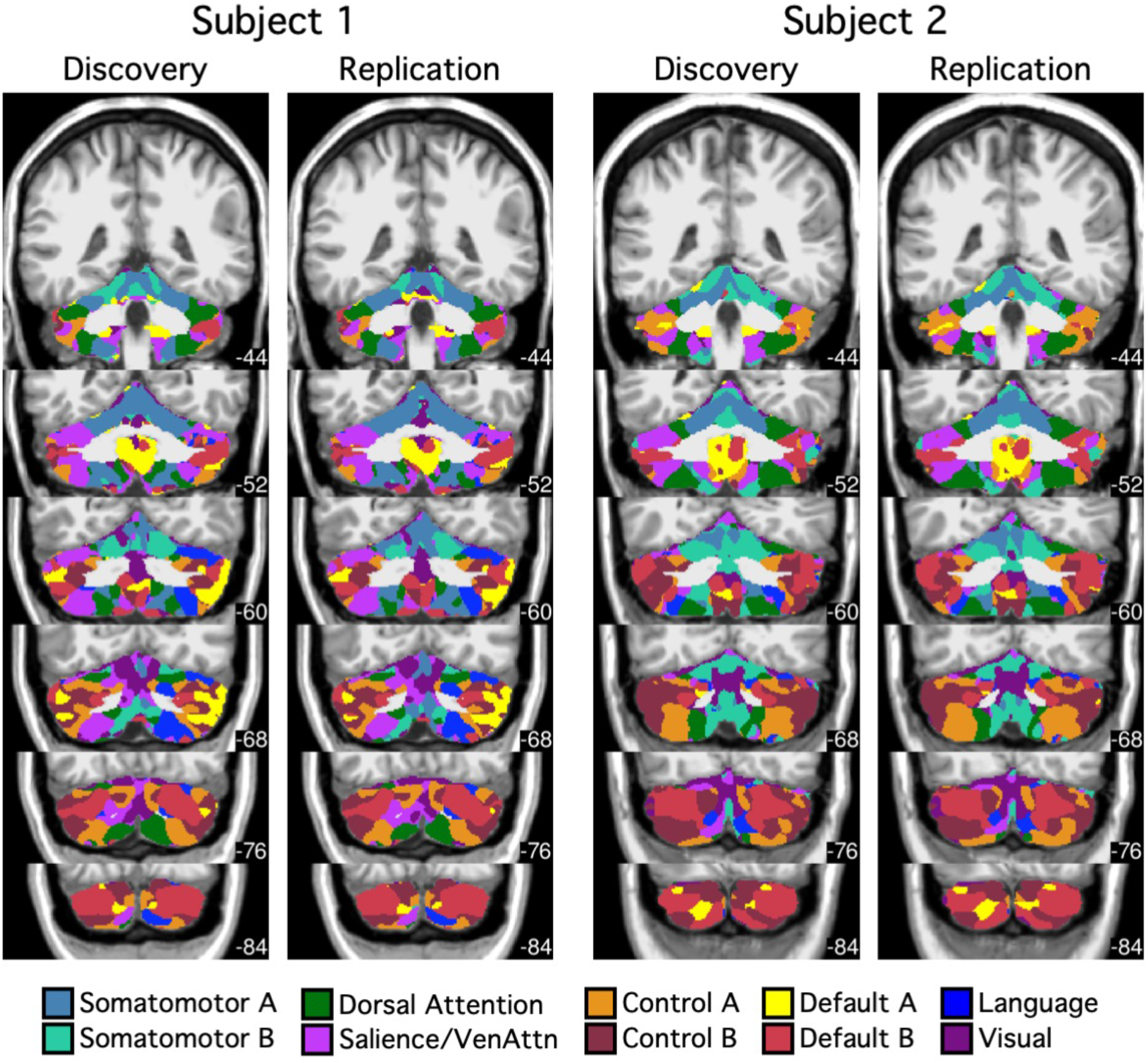
Cerebellar network parcellations were highly reliable across discovery and replication sets within individuals. Individual-specific cerebellar parcellations were estimated using discovery (16 sessions) and replication (15 sessions) datasets independently. Each cerebellar voxel was assigned to the most frequent cortical network among 400 cortical vertices with the strongest correlation (functional connectivity) with the voxel. Individual-specific cerebral parcellations were replicable within subjects. In Subject 1, 83.8% voxels were assigned to the same networks across discovery and replication datasets. In Subject 2, 84.2% voxels were assigned to the same networks across discovery and replication datasets. Networks are colored as labelled in the bottom legend.

### Complete cerebellar maps within individuals

Results above indicate that we were able to derive reliable functional maps within the cerebellum of individuals. To obtain the best estimate of cerebellar maps within each individual, we applied our approach to map the full topography of the cerebellum by using all 62 runs for Subject 1 and all 61 runs for Subject 2. We first estimated the cerebral networks using the MS-HBM approach and then the cerebellar map using a winner-takes-all algorithm.

Figs. 6 and 7 show the individual-specific cerebral cortical parcellations of subjects 1 and 2 respectively. The 10-network cortical parcellations shared some common features as well as clear individual differences across the two subjects. For example, in both participants, default network A and B (yellow/red) were distributed across multiple zones of the cerebral cortex, including the inferior parietal lobule, ventromedial PFC, posterior cingulate cortex, lateral temporal cortex, and other locations, consistent with previous work showing the same broad topography across individuals but with anatomical details varying (Braga et al. 2017; Braga et al. 2019; DiNicola et al. 2020). Similarly, a candidate language network was found in both individuals respecting broad topographic properties that have been well established (e.g., Fedorenko et al. 2012; Mineroff et al. 2018; Braga et al. 2020) but with variability between individuals. In this instance, Subject 1 displayed a more left-lateralized network representation than Subject 2.

**Fig. 6.**
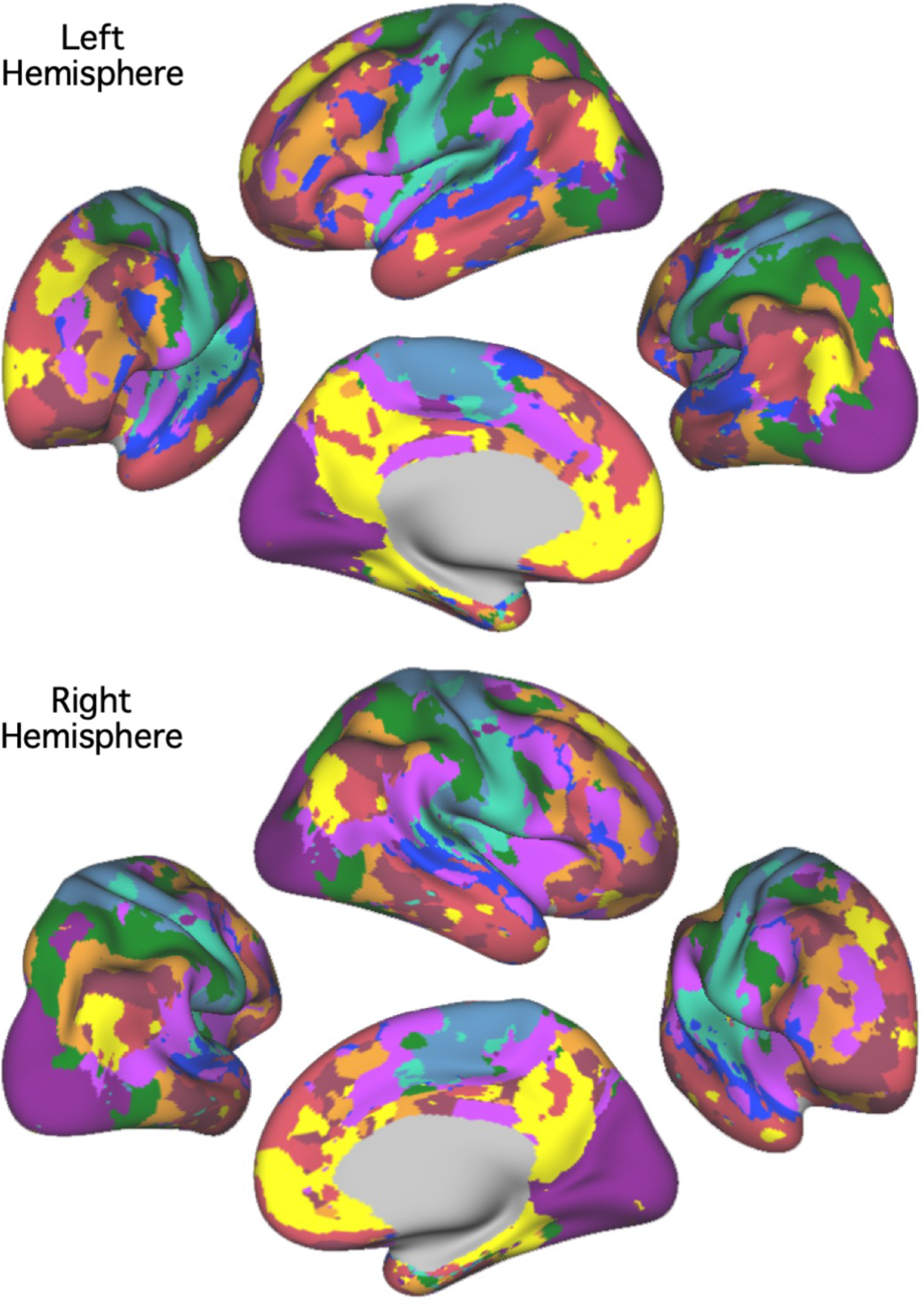
The best estimate of 10-network cerebral cortical parcellation of Subject 1. Multiple views for each hemisphere are provided to show the details of the parcellation. This parcellation represents the best estimate of cortical network organization using the present approach applied to all available data (62 runs from 31 independent sessions). Colors use the network labels as shown in Fig. 4. Note the presence of juxtaposed interdigitated networks in high-order association cortex that include default network A (yellow) and default network B (red), a candidate language network (blue), and cognitive control networks (orange and brown).

**Fig. 7.**
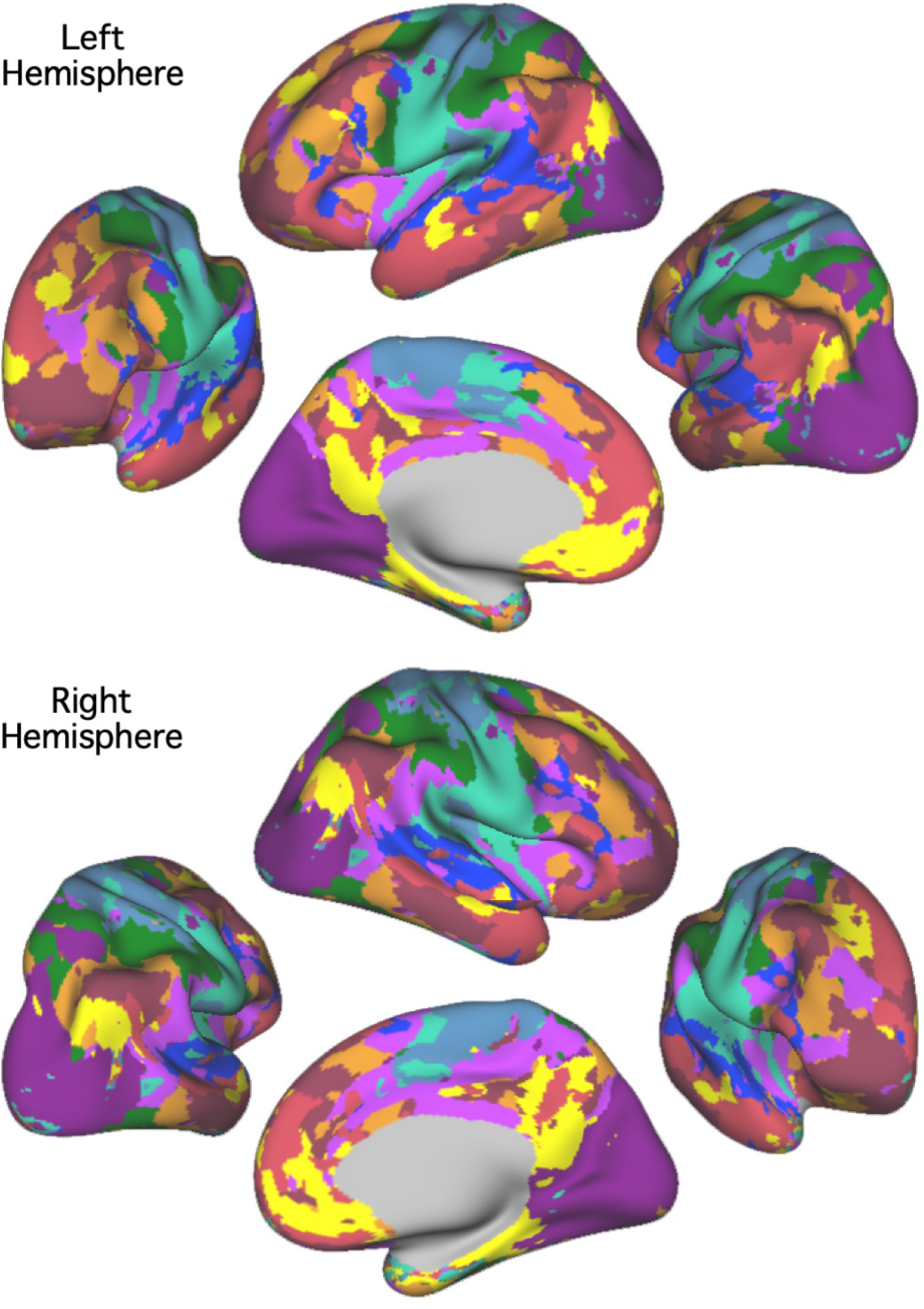
The best estimate of 10-network cerebral cortical parcellation of Subject 2. Multiple views for each hemisphere are provided to show the details of the parcellation. This parcellation represents the best estimate of cortical network organization using the present approach applied to all available data (61 runs from 31 independent sessions). Colors use the network labels as shown in Fig. 4. Note the presence of juxtaposed interdigitated networks in high-order association cortex that include default network A (yellow) and default network B (red), a candidate language network (blue), and cognitive control networks (orange and brown).

Figs. 8 and 9 show the individual-specific cerebellar parcellations of Subjects 1 and 2 in the volume and Fig. 10 shows the parcellations on the cerebellar surface. Like the cerebral cortex, the cerebellar parcellations of the two subjects displayed common as well as individual-specific features. First, for both subjects, the somatomotor regions in the anterior and posterior lobes were mapped to somatomotor networks in lobule IV-VI and lobule VIIIb. More specifically, foot and hand regions were assigned to two fractionated networks, provisionally labeled somatomotor network A (blue) and somatomotor network B (aqua). Of note the two separate somatomotor network assignments capture features of the body map topography observed in targeted analyses (e.g., Fig. 3) but not all of the organization. Second, for both subjects, the majority of the cerebellum was mapped to cerebral association networks, including Crus I/II, lobule IX, X, and part of lobule VI, VIIb, VIIIa. However, the size and location of the association networks were different between the two subjects. For example, similar to the cerebral cortex, the language network representation within the cerebellum was more lateralized in Subject 1 than in Subject 2. Thus, while the language network had generally similar organizational features between the two subjects (note the clear parallels between the multiple distributed representations in the right cerebellar hemisphere) there were also differences that suggest individual variations in exact spatial locations as well as laterality.

**Fig. 8.**
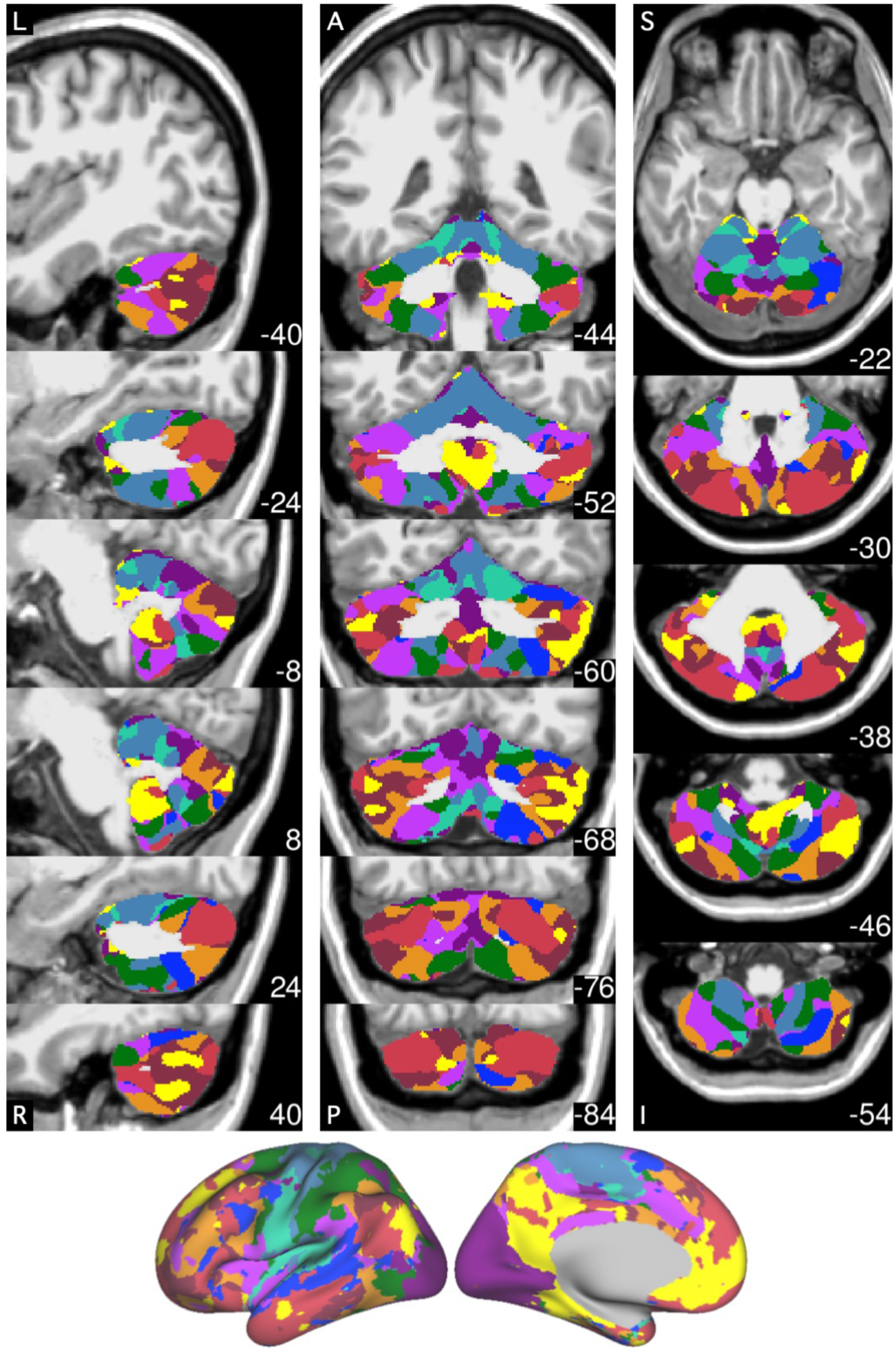
The best estimate of 10-network cerebellar parcellation of Subject 1. Each cerebellar voxel was assigned to the most frequent cortical network among the 400 cortical vertices with the strongest correlation (functional connectivity) with the voxel. The 10 cerebral networks of Subject 1 are shown at the bottom for reference. Colors use the network labels as shown in Fig. 5. The three sections display sagittal (left), coronal (middle), and axial (right) views. L: left; R: right; A: anterior; P: posterior; S: superior; and I: inferior. The left hemisphere is displayed on the left. Coordinates at the bottom right of each panel indicate the section level in MNI152 space. Each network identified in the cerebral cortex has multiple representations in the cerebellum. The organization is broadly symmetric between the cerebellar hemispheres, with asymmetries paralleling the cerebral network asymmetries. For example, the language network (blue) shows a markedly expanded representation in the right hemisphere consistent with the leftward asymmetry in the cerebral cortex.

**Fig. 9.**
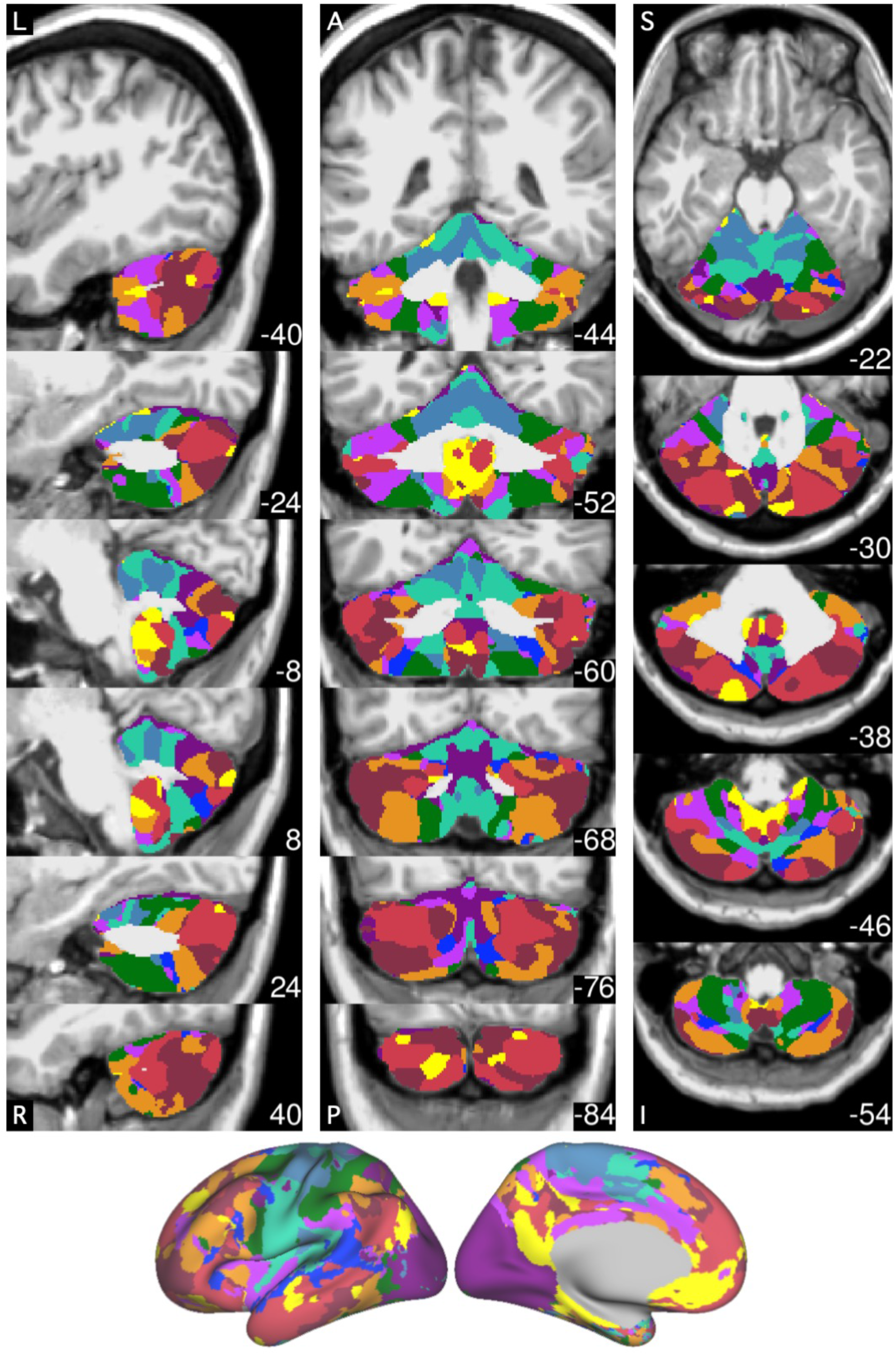
The best estimate of 10-network cerebellar parcellation of Subject 2. Each cerebellar voxel was assigned to the most frequent cortical network among the 400 cortical vertices with the strongest correlation (functional connectivity) with the voxel. The 10 cerebral networks of Subject 2 are shown at the bottom for reference. Colors use the network labels as shown in Fig. 5. The three sections display sagittal (left), coronal (middle), and axial (right) views. L: left; R: right; A: anterior; P: posterior; S: superior; and I: inferior. The left hemisphere is displayed on the left. Coordinates at the bottom right of each panel indicate the section level in MNI152 space. Each network identified in the cerebral cortex has multiple representations in the cerebellum.

**Fig. 10.**
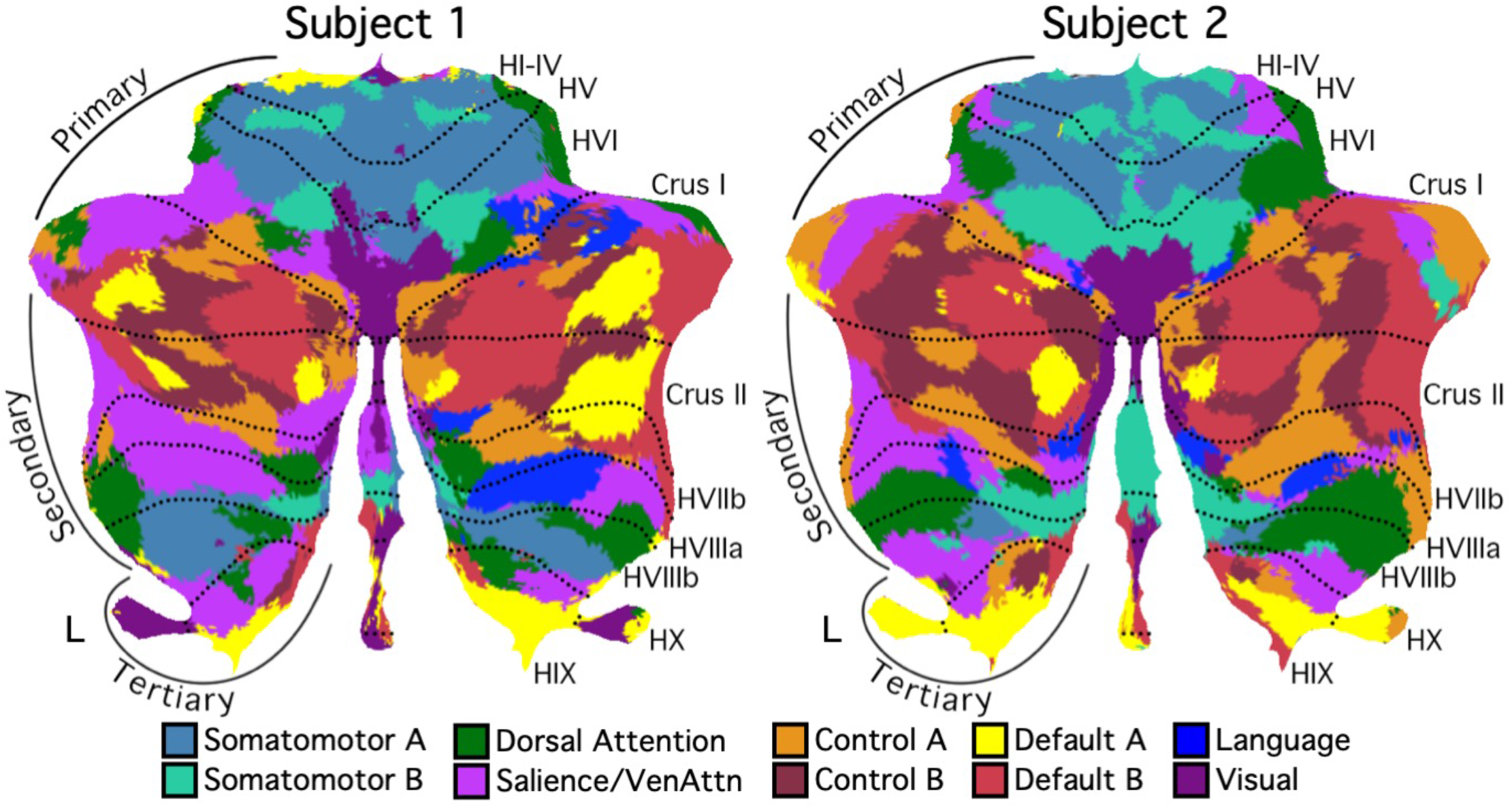
Cerebellar network parcellations within individuals are shown on flatmaps. The individual-specific parcellations were generated using all sessions for each subject projected using the SUIT toolbox (Diedrichsen et al. 2015). Dotted lines indicate lobular boundaries. L indicates left cerebellar hemisphere. Different lobules are marked on the right side. Networks are colored as labelled at bottom. Despite the complex topography, the cerebellum contained three representations of the cerebral cortex labelled as the putative Primary, Secondary, and Tertiary maps. The Primary map begins with the anterior lobe somatomotor representation, passes through dorsal attention and salience/ventral attention networks, and ends in the apex association networks centered at the Crus I/II border, including clear representation of default network A and B. The Secondary map then progresses in reverse through to a second Somatomotor representation in the posterior lobe (within HVIIIb). Evidence for the Tertiary map is the final representation of networks between HVIIIb and HIX that possesses representation of default network A and B in HIX.

### Individual cerebellar topography includes multiple representations of the cerebrum

Our previous explorations suggested that at the group-level, the cerebellum contains three representations of the cerebral cortex (Buckner et al. 2011; see also Guell et al. 2018a). Despite the complex topography of individual-specific cerebellar parcellations, the three representations could also be observed in both individual subjects (Figs. 10 and 11). Fig. 11 shows the three representations in different sagittal sections. The three representations are also marked on the left side of the flatmaps in Fig. 10.

**Fig. 11.**
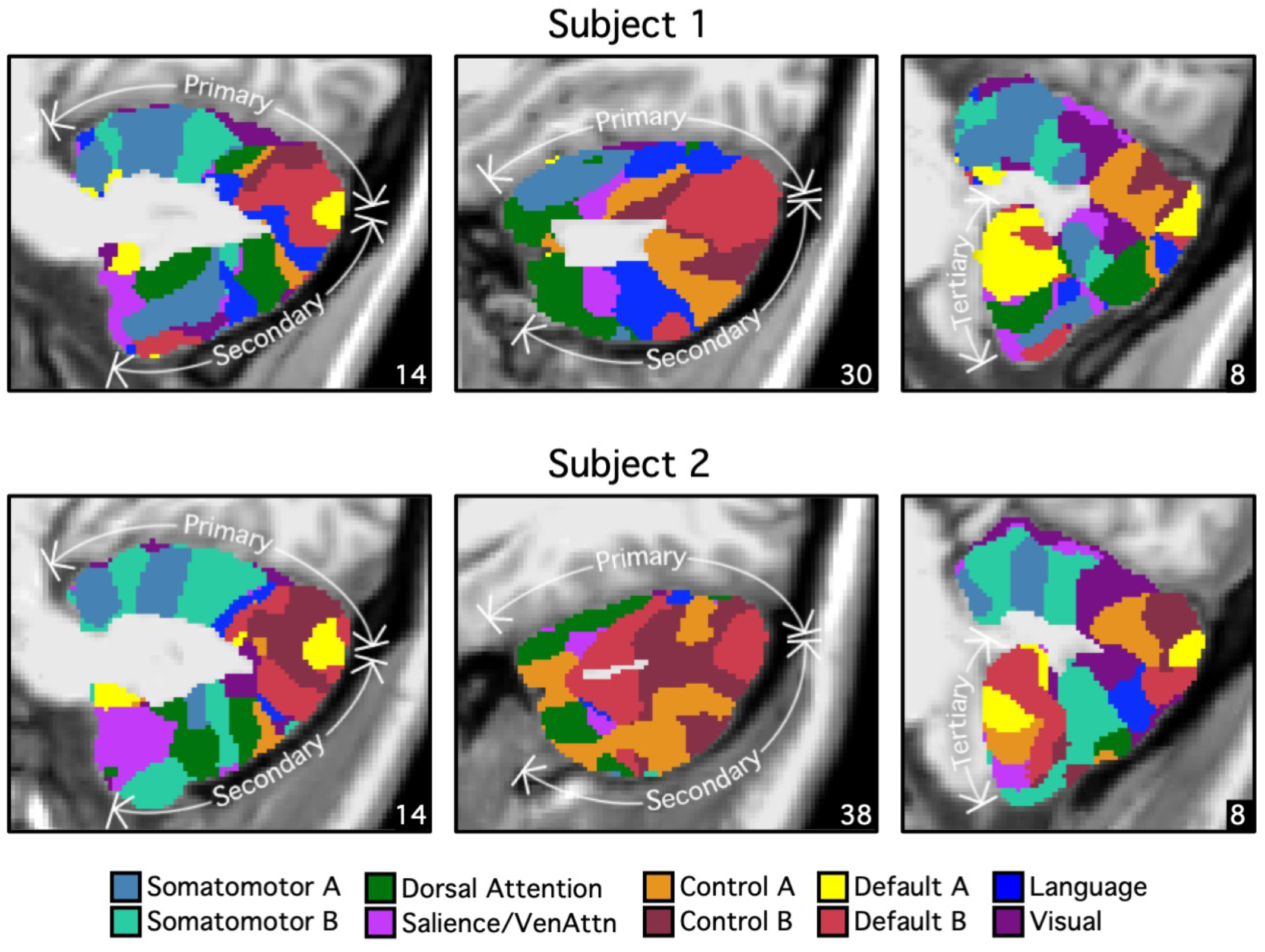
Cerebellar network parcellations within individuals shown in the volume to reveal multiple repeating maps. The topographic ordering of the cerebral networks from Fig. 10 is illustrated for 3 sagittal sections of the right cerebellum. Three distinct representations are observed for both subjects, labeled as Primary, Secondary, and Tertiary. Each map is a roughly duplicated ordering of the adjacent map (with some variation).

The primary map began with the somatomotor networks at the anterior lobe and ended with the default network A and B at Crus I/II. The remaining association networks were sandwiched in between the somatomotor and default network A and B. Association networks that are more involved in sensory perception (i.e., dorsal and ventral attention networks) were positioned adjacent to the somatomotor networks, while association networks involved in cognitive control and language were positioned adjacent to the default network A and B. Default network A was juxtaposed to default network B in multiple locations in both individuals.

The secondary map began with default network A and B at Crus I/II and ended with somatomotor networks at lobule VIIIb. The ordering of the networks in the primary and secondary maps were basically in an inverse order but with some variation. The tertiary map was present in lobules VIIb, IX, and X. Not all association networks were independently observed in the smaller tertiary map, but default network A and B were particularly robust and distinct (Fig. 12).

**Fig. 12.**
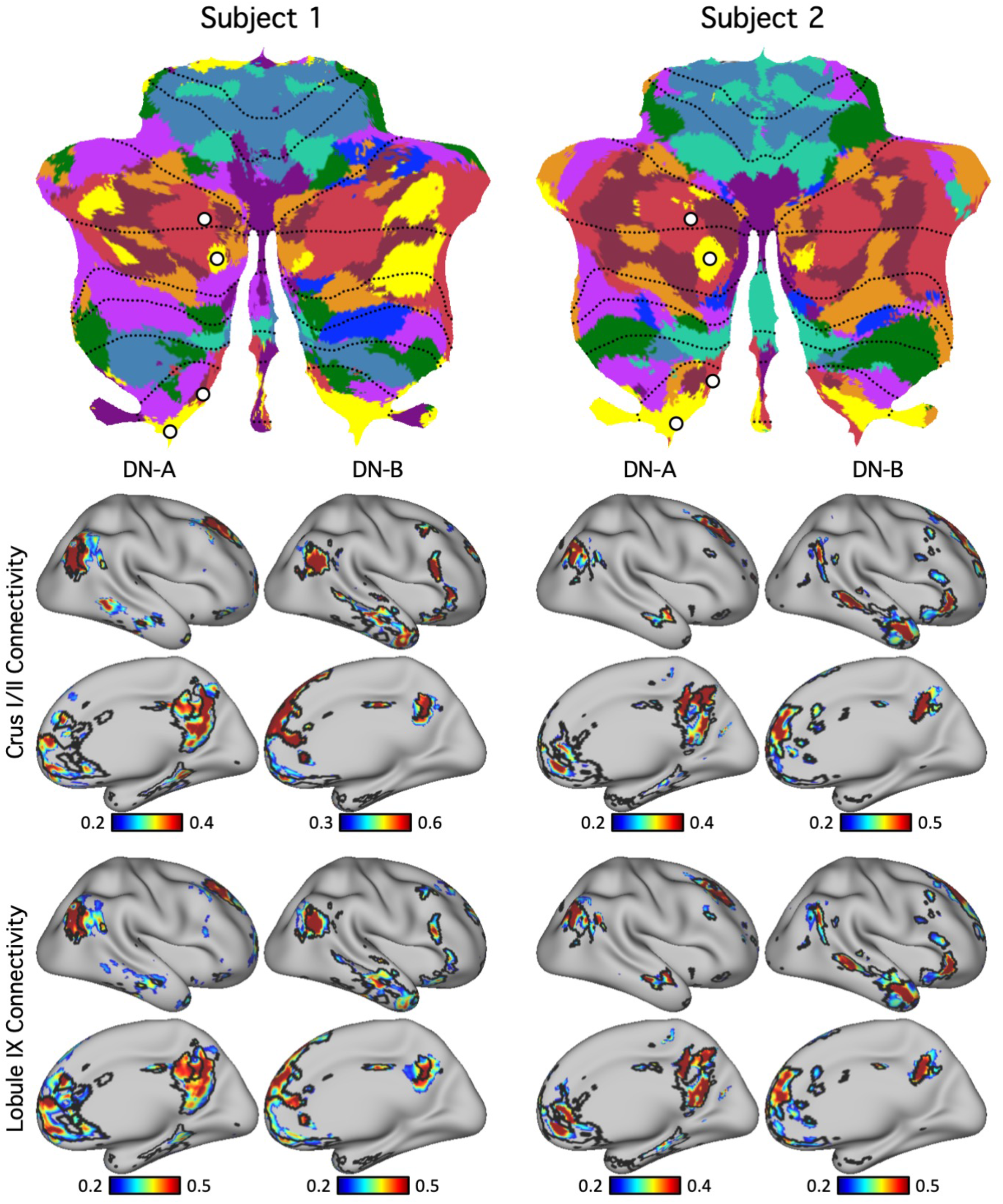
Evidence for specificity of default network A and B. (Top panel) Seed regions from default network A and B were selected within the discovery set (plotted as white circles). Note two pairs of seed regions were selected, centered on adjacent default network A and B representations separately for the Crus I/II representation and the spatially distant HIX representation. This allowed the network topography of each separated cerebellar map to be examined in the cerebral cortex. (Bottom panel) Functional connectivity maps from the cerebellar seed regions were estimated for the cerebral cortex using the replication set. Black lines indicate boundaries of individual-specific default network A or B from the original cerebral parcellation (Fig. 4, replication set). Each pair of juxtaposed cerebellar seed regions yielded the full, distributed cerebral networks associated with the separate default network A or B; seed regions from the same networks in Crus I/II and Lobule IX exhibited highly similar functional connectivity patterns. These results illustrate remarkable specificity of the cerebellar parcellations.

Thus, the broad hierarchical ordering that has been previously highlighted in relation to cerebral network organization (Margulies et al. 2016; Braga and Buckner 2017) reveals a parallel macroscale organization in the cerebellum with a primary, secondary, and likely tertiary map of the network set. However, the exact details of the adjacencies also show variability between subjects and complex juxtapositions, especially among the higher-order association networks. This is evident near the Crus I and II border. Specifically, in both individuals, the default network A and B lie side by side adjacent to other higher-order association networks including control networks A and B, and the language network. This complex topography in the cerebellum is reminiscent of the complex interdigitations of networks that exist in the multiple association zones of the cerebral cortex (e.g., Fedorenko et al. 2012; Braga and Buckner 2017; Gordon et al. 2017; Braga et al. 2019; Gordon et al. 2020). Despite this complex topography between the higher-order association networks, the broad hierarchical ordering was largely preserved with the default network A and B at the apex in both subjects. Representations of the dorsal attention network were distant from the apex networks in both the anterior and posterior lobe, and the somatomotor representations were the farthest. One unexpected exception to this hierarchical arrangement, concerning a novel representation of early visual cortex in the cerebellum, will be reported in detail in a later section.

### Seed-based functional connectivity demonstrates specificity of cerebrocerebellar circuits

The detailed and complex organization of the cerebellum described above relies on several assumptions including the use of a winner-take-all parcellation algorithm that could give an appearance of specificity with non-specific features and spatial blurring deemphasized. The methods also assume there is a direct relationship between the cerebral networks and the cerebellar networks. If a cerebellar region exhibited a different connectivity pattern in the cerebral cortex, it would still be assigned to the best-fitting network. For this reason, like in our original efforts to identify network organization in averaged subject groups (Yeo et al. 2011; Buckner et al. 2011), we examined cerebellar organization using an independent method that made few assumptions about the spatial details of cerebral network organization. The basic idea here is that all analysis methods are biased and true features of organization should be robust to multiple, distinct analysis strategies.

For these analyses, we selected seed regions in the cerebellum using the discovery set and visualized their functional connectivity patterns in the cerebral cortex in the replication data set for each individual. Across seed pairs we asked a series of questions to test whether dissociations existed between nearby cerebellar regions as well as to test whether distributed regions in the multiple cerebellar representations link to the same cerebral network as hypothesized.

To confirm default network representation within the cerebellum, we selected four cerebellar seed regions in each subject. To capture primary/secondary representations, two seed regions were selected in Crus I/II: one in default network A and one in default network B. To capture tertiary representations, two seed regions were selected in lobule IX: one in default network A and one in default network B. In both subjects, the resulting cerebrocerebellar functional connectivity maps were in strong agreement with individual-specific default network boundaries in the cerebral cortex (Fig. 12). This is a critical result that simultaneously shows the spatial specificity of the distinct default network A and B representations and that the specific representations in spatially discontinuous zones of the cerebellum converge on the same cerebral networks.

Similarly, the functional connectivity patterns of seed regions in the cerebellar language representations showed agreement with individual-specific language boundaries in the cerebral cortices of both subjects (Fig. 13). While both subjects exhibited three representations of the cerebral cortex within the cerebellum, there were individual differences in the network topography. Of interest, the differences suggesting idiosyncratic spatial positions of the language network based on the winner-take-all parcellation (the multiple blue zones in the flat maps) were confirmed by seed-based functional connectivity. Each of the three representations of the language network in the right cerebellar hemisphere in each individual yielded a complete map of the spatially-specific language network in each subject (Fig. 13).

**Fig. 13.**
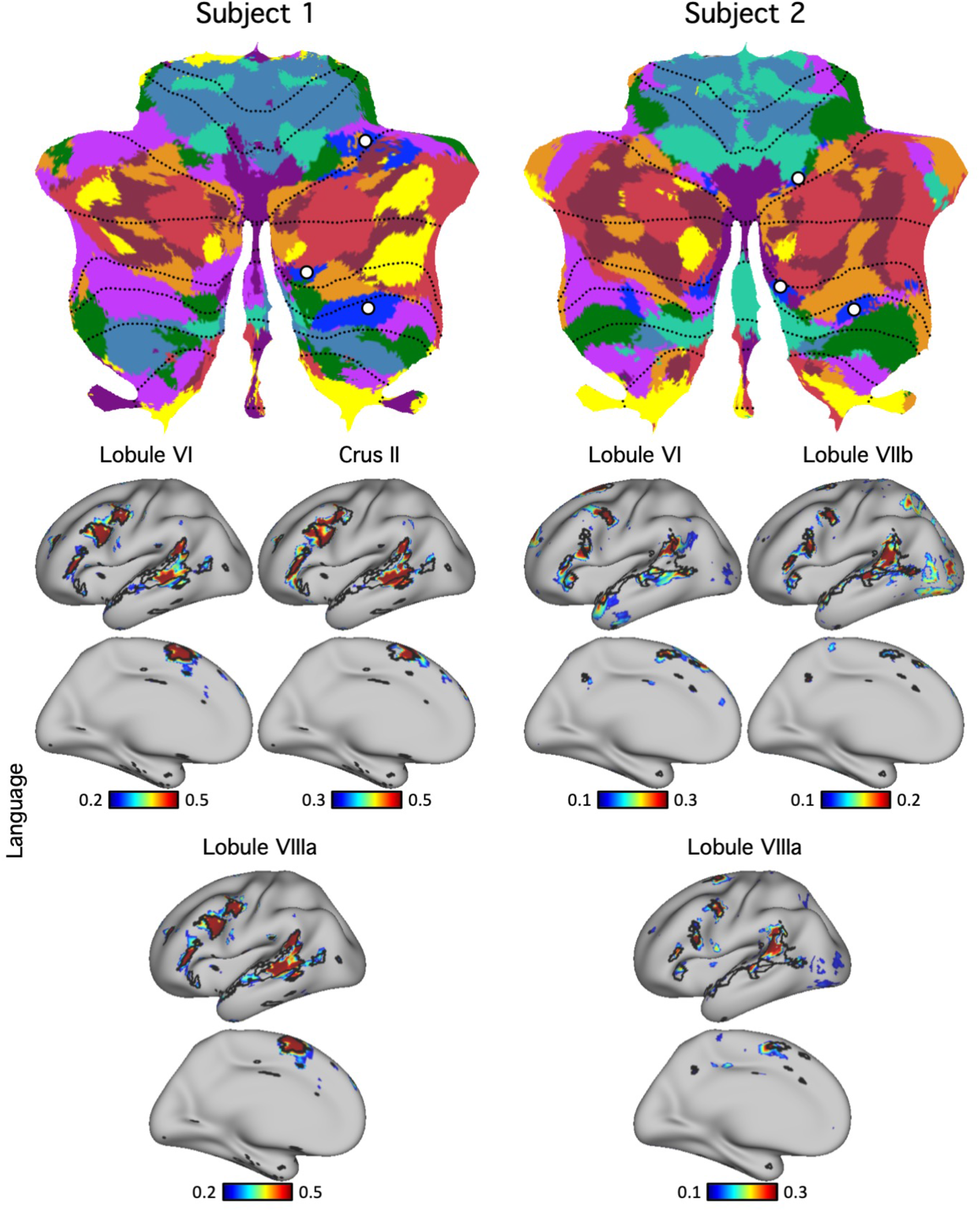
Evidence for specificity of the language network. (Top panel) Cerebellar seed regions from the language network were selected using the discovery set. Three separate seed regions were selected within each individual that included representations in the spatially distant HVI and HVIIIa regions. (Bottom panel) Functional connectivity maps of these seed regions were estimated using the replication set. Black lines indicate boundaries of the individual-specific language network and highlight agreement between the network estimates from each of the three cerebellar representations.

Distinctions between networks were also confirmed. As an example, Fig. 14 shows the functional connectivity patterns of two pairs of cerebellar seed regions assigned to different networks by the winner-takes-all algorithm. One seed region was located in the language network in Subject 1 and default network B in Subject 2. The second seed region was located in default network A in Subject 1 and control network B in Subject 2. The functional connectivity patterns of these seed regions agreed well with individual-specific cerebral cortical network boundaries (Fig. 14).

**Fig. 14.**
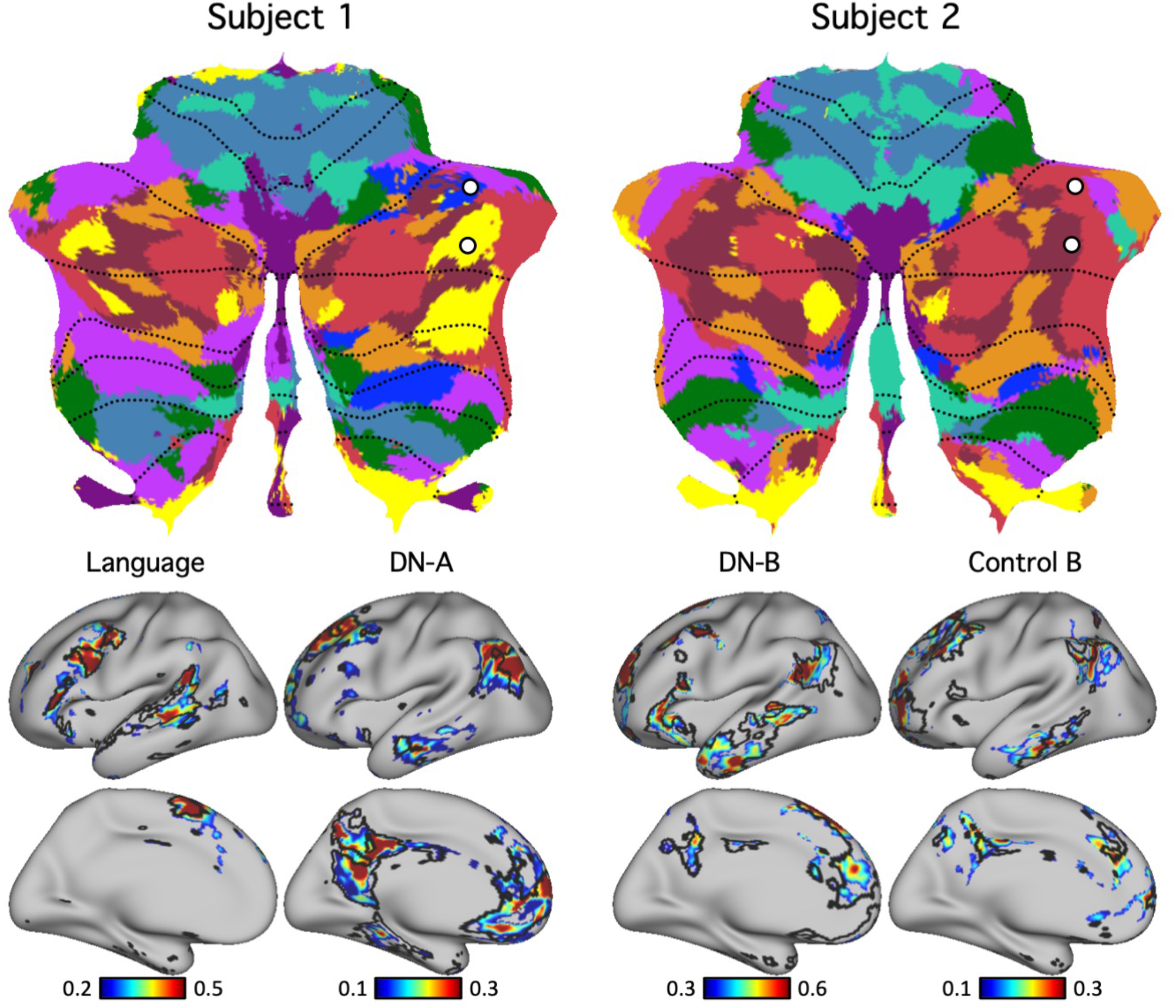
Differences in cerebellar topography between individuals were supported by seed-based functional connectivity. (Top panel) Pairs of seed regions were selected in the discovery dataset to highlight topographic differences between the two individuals. To make this point most clearly, seed regions were selected for the same MNI coordinates in both subjects, with the corresponding networks they aligned to differing between subjects. The first cerebellar seed region was within the language network in Subject 1 and within the default network B in Subject 2. The second cerebellar seed region was within default network A in Subject 1 and within control network B in Subject 2. (Bottom panel) Functional connectivity maps of these seed regions were estimated using the replication sets. Black lines indicate boundaries of corresponding individual-specific networks and highlight strong agreement with functional connectivity maps. These results demonstrate how zones of the cerebellum that are located in similar volumetric positions between individuals can correspond to starkly different functional networks.

An additional feature of the winner-take-all parcellation was examined that surrounded seemingly ‘ectopic’ network representations. In Subject 2, there was a zone in the lateral right cerebellar hemisphere assigned to somatomotor cortex, surrounded by zones of higher-order association cortex. An intriguing possibility is that such a zone might emerge as a local ectopic specialized region of the cerebellum with a discontinuity from neighboring networks, at least as conceived in a hierarchical organizational framework. We did not find support for this possibility. Fig. 15 shows a seed region placed in the candidate ectopic somatomotor representation. Unlike the dissociations described above, where clear spatially-specific network patterns emerged, here a noisy cerebral network pattern emerged. These discrepancies might be due to low signal-to-noise or biological origins that our approach was unable to capture.

**Fig. 15.**
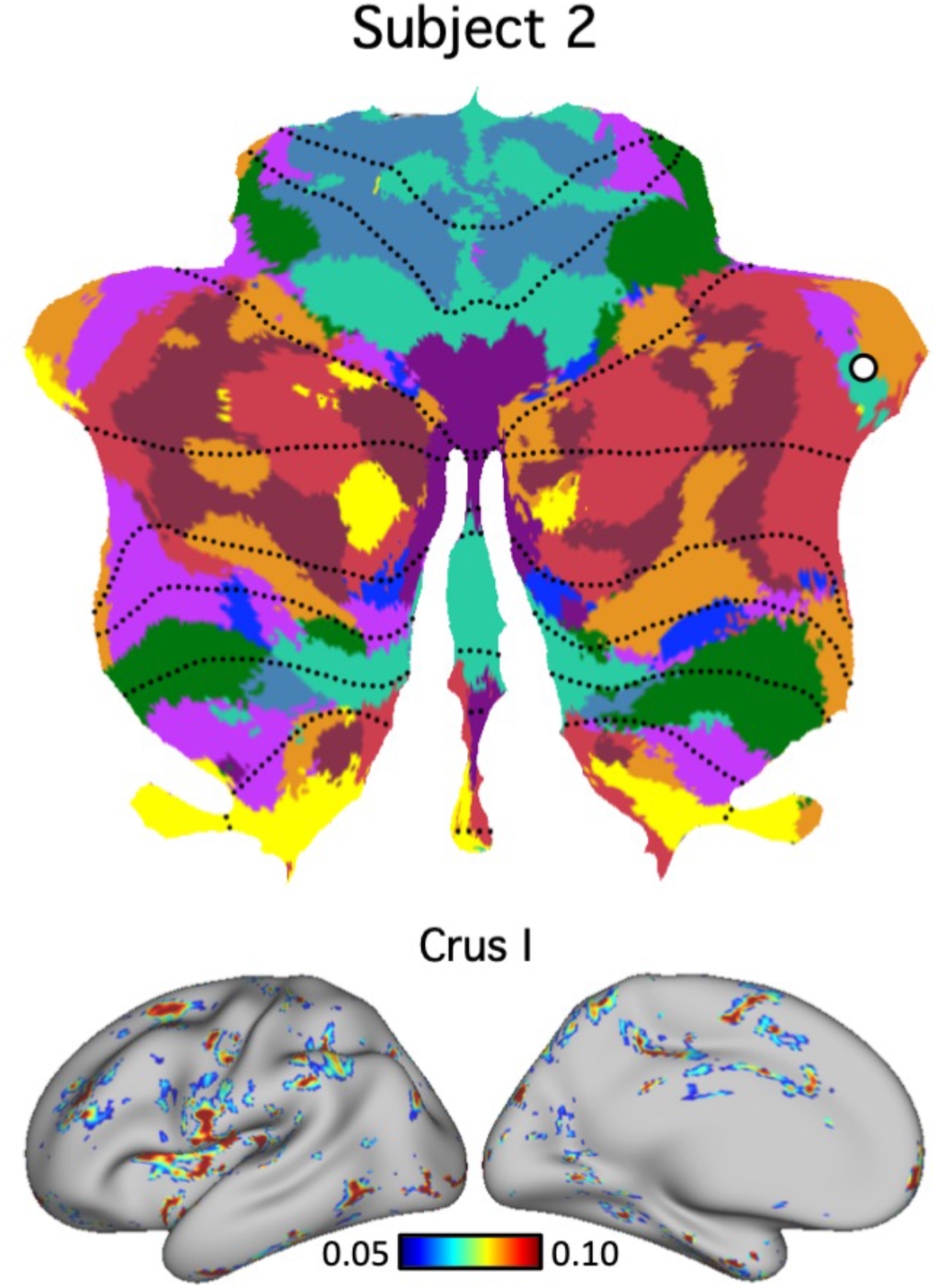
A failed test of an ectopic cerebellar functional zone. (Top panel) A cerebellar seed region was selected using the discovery set in a region that showed an ectopic network assignment. The region was assigned to a somatomotor network in a zone of the lateral hemisphere near Crus I that is otherwise surrounded by higher-order association networks. (Bottom panel) The functional connectivity map of the seed region estimated using the replication set reveals a distributed, noisy pattern. This pattern is not easily interpreted and may either represent an alternative biological arrangement or experimental noise that we presently do not understand. Failures of this type were rare.

### Visual representation within the cerebellum of individuals

Analyses of the cerebellum in humans and non-human primates have suggested that early retinotopic visual areas do not have a representation in the cerebellum, although recent analysis of high-resolution human data have suggested candidate zones that respond to retinotopic visual stimuli (van Es et al. 2019). The cerebellar maps in both individuals analyzed here support the existence of cerebellar representation of early retinotopic visual cortex. This visual representation was mainly present in the vermis with some extension to other lobules (mainly lobule VI) as can be seen in both subjects in the volume (Figs. 8 and 9) and flatmap (Fig. 10), corresponding near to the oculomotor vermis (OMV) representation of van Es et al. (2019). Functional connectivity maps of cerebellar seed regions were in agreement that these regions coupled to early visual cortex in both individual subjects, and further that they could be dissociated from higher-order visual representations linked to the dorsal attention network (Fig. 16).

**Fig. 16.**
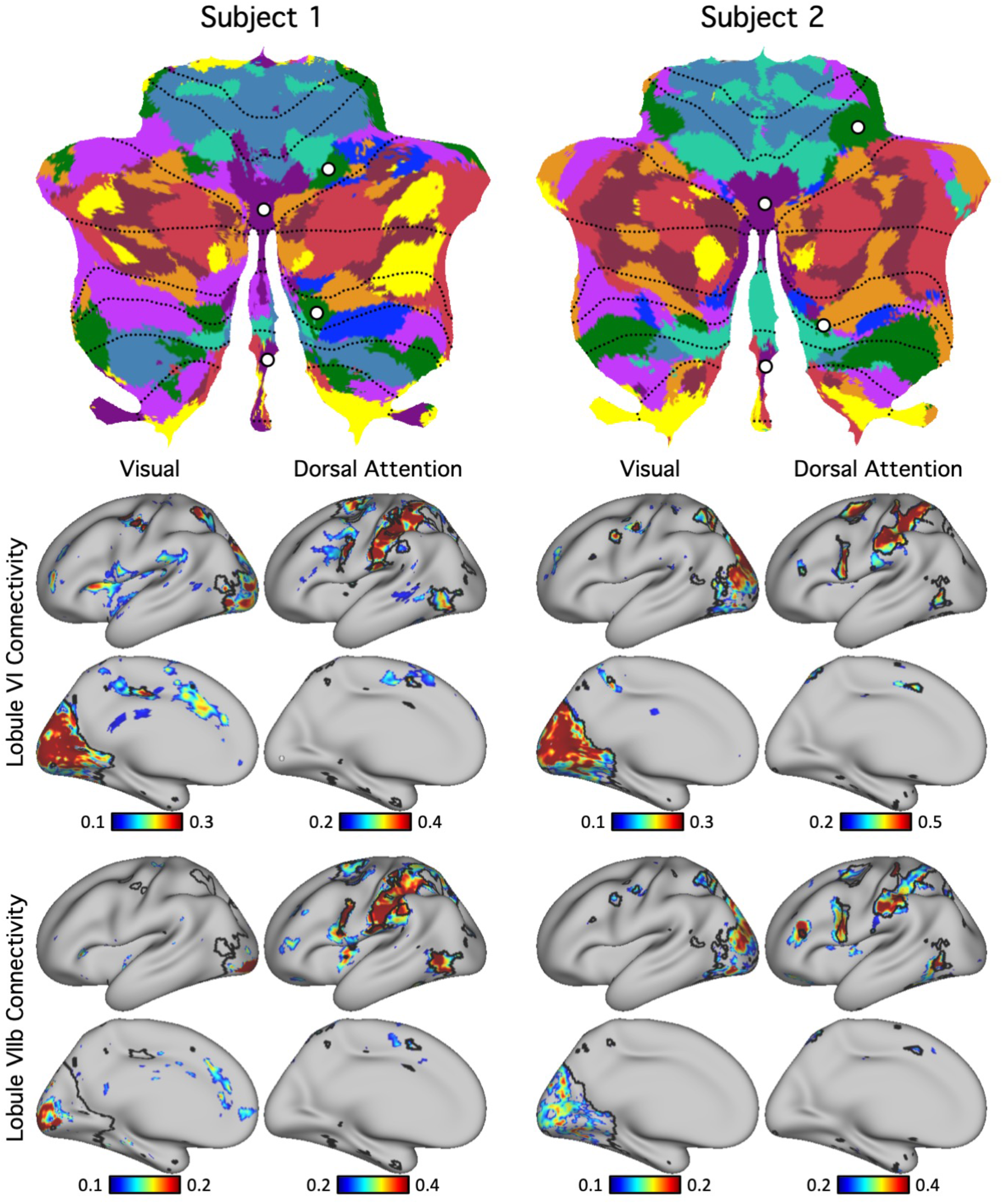
Evidence for specificity of a visual network representation in the cerebellum. (Top panel) Cerebellar seed regions were selected using the discovery set within a region of the vermis assigned to the visual network (purple) and a spatially separate region assigned to the dorsal attention network. (Bottom panel) Functional connectivity maps of these seed regions were estimated using the replication set. Black lines indicate boundaries of individual-specific visual and dorsal attention networks and highlight agreement with functional connectivity maps. The functional connectivity maps support the existence of early visual representation within the vermis of the cerebellum. Of note, the cerebral cortical regions coupled to the cerebellar vermis included early retinotopic visual cortex at or near V1.

We note that a small visual region inside the vermis was also present in our previous group-level cerebellar parcellation (Fig. 17; Buckner et al. 2011). Red voxels in the middle of the vermis were assigned to visual network B (Yeo et al. 2011). Comparing the group-level parcellation (Fig. 17) with our individual-specific parcellations (Figs. 8 to 10), the visual representation within the group-level parcellation was limited to the vermis.

**Fig. 17.**
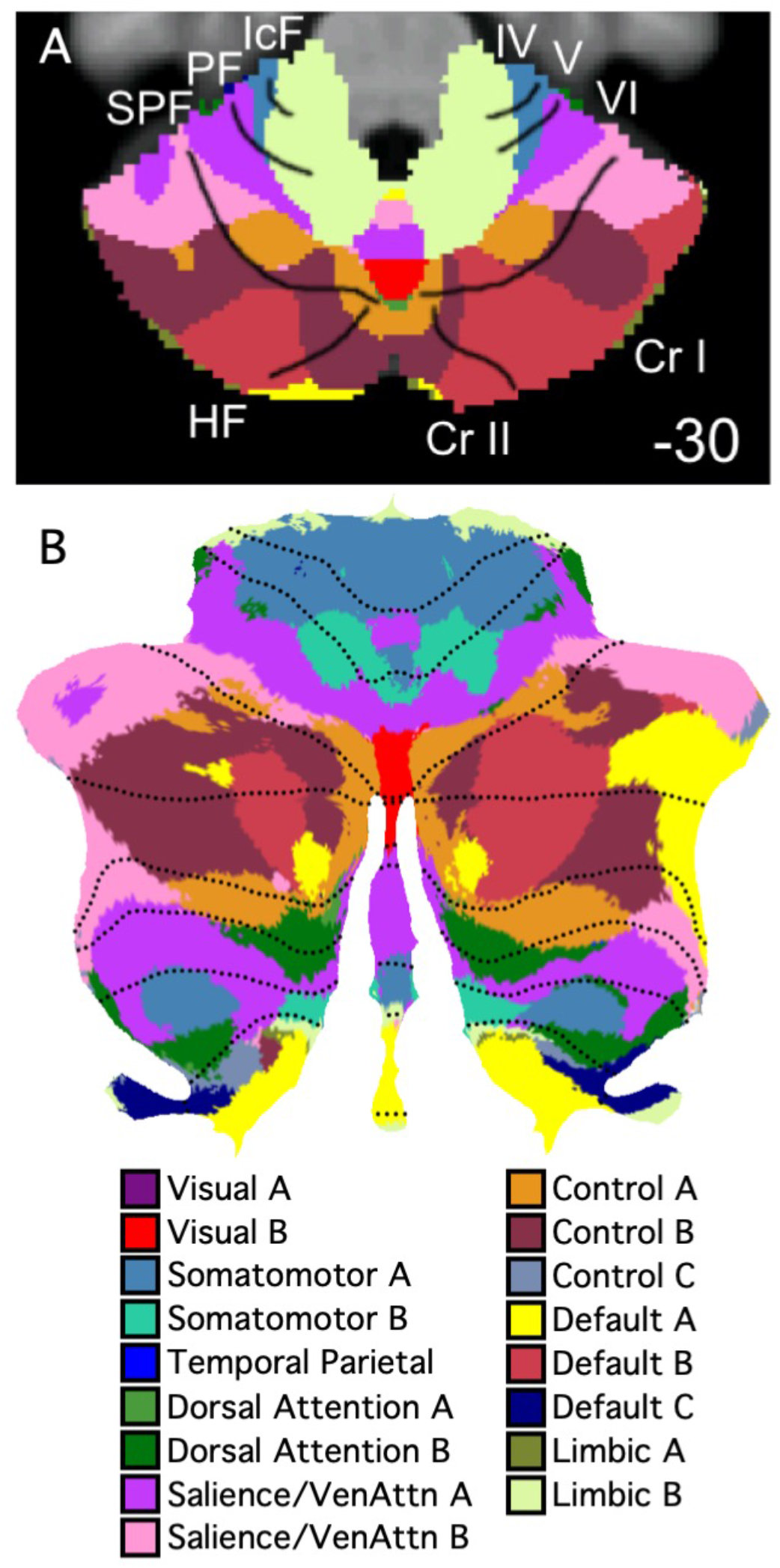
Evidence for visual network representation in group-averaged data. Group-level 17-network cerebellar parcellation from Buckner et al. (2011) also revealed visual representation (visual B in red) in the vermis. (A) An axial section shows visual network B in the vermis. (B) The same data projected to a flatmap. The visual region appeared in roughly the same location in both the group-level and individual-specific cerebellar parcellations (see Fig. 16).

Further exploration of seed regions within the vermis suggested that different seed locations exhibited preferential functional connectivity with central and peripheral locations of visual cerebral cortex (Fig. 18). To ensure that the visual representation was not an artifact of signal blurring between the cerebral cortex and cerebellum, we selected seed regions from the visual cerebral cortices (left panel in Fig. 19) and computed their functional connectivity in the volume (right panel in Fig. 19). The bright correlated regions within the cerebellum (white circles in Fig. 19) were clearly distant from the occipital lobe for both subjects, suggesting that the visual representation within the vermis was unlikely to be an artifact of signal blurring.

**Fig. 18.**
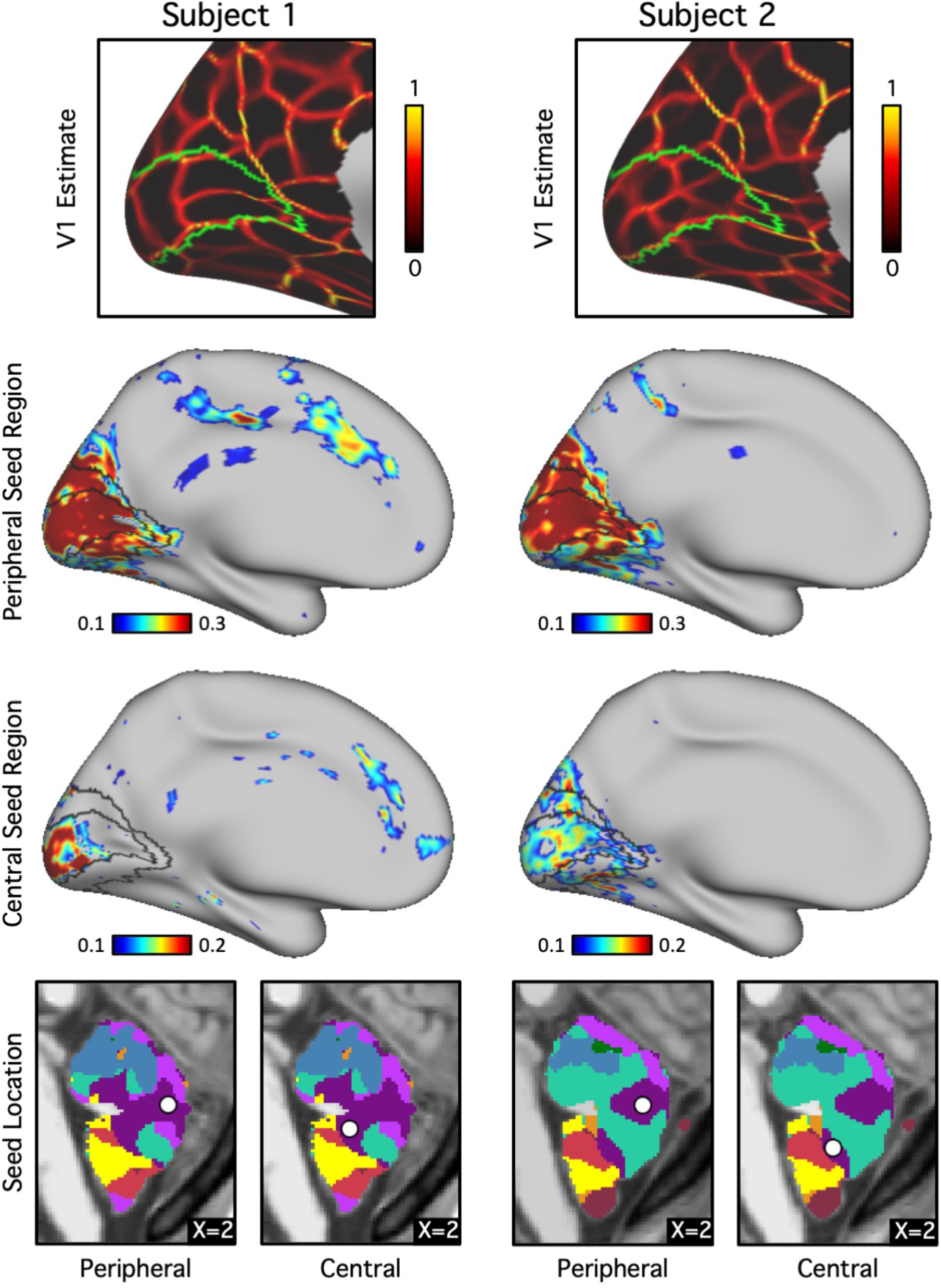
Evidence for central and peripheral visual representations coupled to primary visual cortex within the cerebellum. (Top row) V1 is estimated in each individual based on projected histology (green line) and within-individual functional connectivity gradients (red-yellow lines). Histological estimates are derived from the Juelich atlas in FreeSurfer (Amunts et al. 2000; Fischl et al. 2008). The functional connectivity gradient maps (Laumann et al. 2015; Gordon et al. 2016) show sharp transitions in functional connectivity patterns across the cerebral cortex and align with the histological V1 boundaries. (Middle rows) Cerebral functional connectivity maps are displayed from seed regions placed in the visual zone of the cerebellar vermis. Histological V1 and V2 boundaries from the Juelich atlas in FreeSurfer are overlaid in black. Note that functional connectivity patterns strongly overlap with the estimate of primary visual cortex (V1). Separate regions are associated more with the peripheral and central visual zones, particularly evident for Subject 1. (Bottom row) The locations of the cerebellar seed regions are displayed in sagittal sections.

**Fig. 19.**
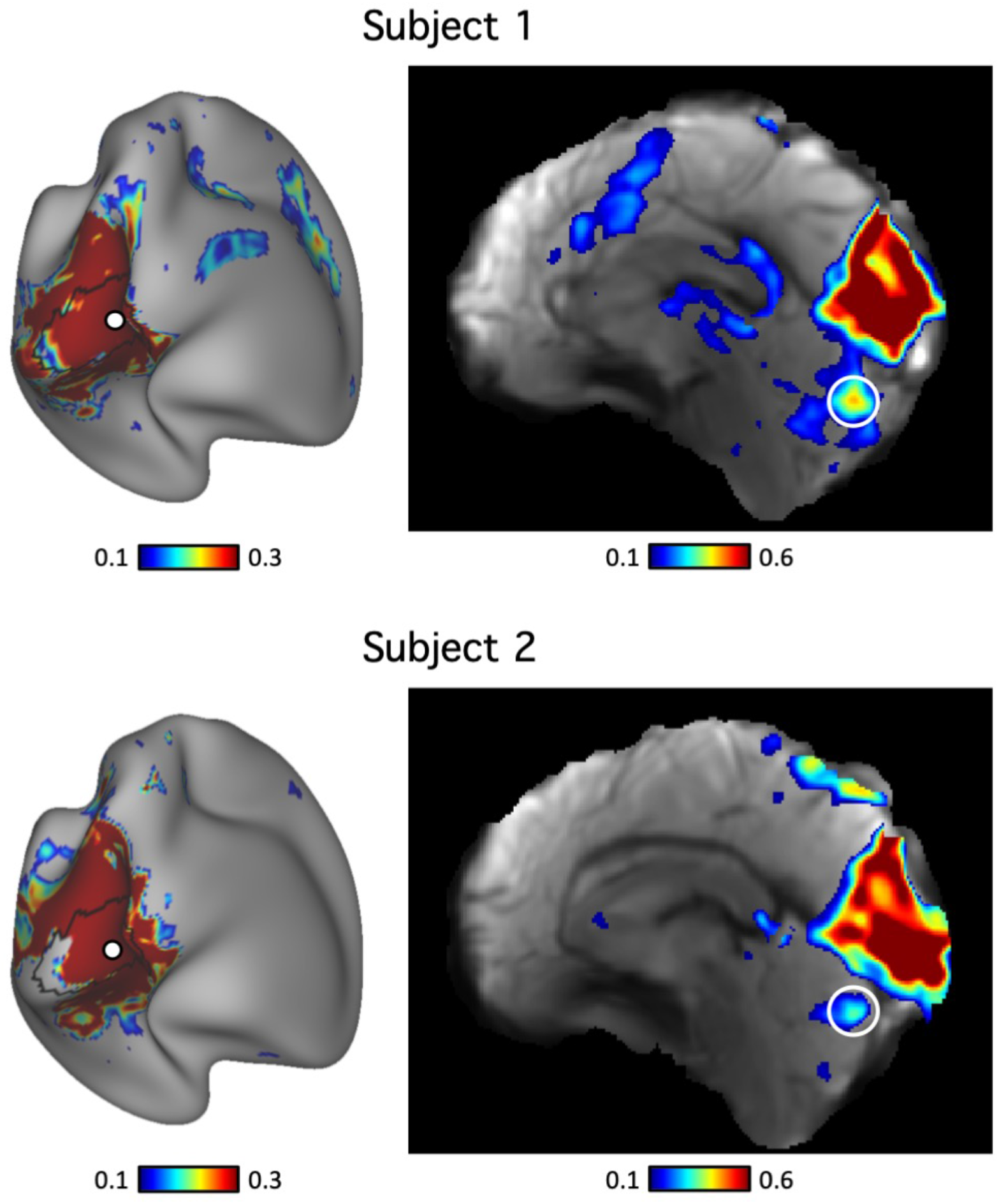
Primary visual cortex connectivity reveals selective representation in the cerebellum within individuals. Seed regions were selected in the visual cerebral cortex in the discovery set. The seeds regions are shown as white circles in the left panels. Functional connectivity maps of the seed regions were computed in the volume using the replication dataset and shown in the right panels. Cerebellar islands (white circles) show high connectivity with cortical visual regions that are separate from the correlation within and around the seed region in the cerebral cortex (the large region above), suggesting that the visual representation within the vermis is unlikely to be the result of signal leakage from the occipital lobe.

### The cerebellum is proportionately mapped to the cerebral cortex

The final analysis explored the relationship between the extent of each network representation in the cerebral cortex to that of its representation in the cerebellum (paralleling Buckner et al. 2011 and Marek et al. 2018). For this analysis, the networks were analyzed separately for the left-and right-hemispheres so as to allow lateralized networks to reveal their organization fully. Fig. 20 shows the relationship between the extent of cerebral and contra-lateral cerebellar cortices assigned to distinct functional networks.

**Fig. 20.**
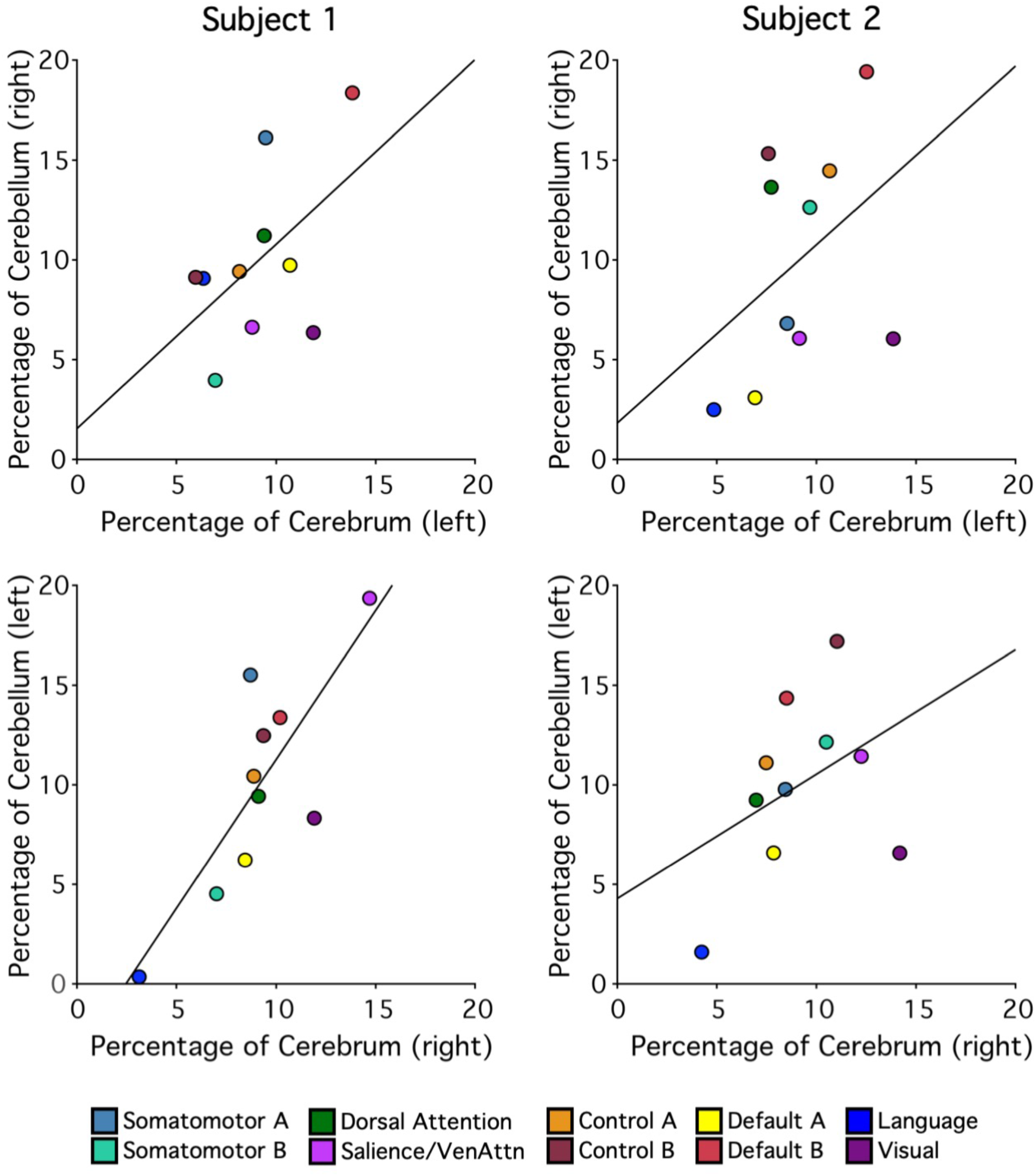
Relationship between the extent of cerebral and contra-lateral cerebellar cortices assigned to distinct functional networks. Each colored circle represents a different network as labelled at bottom. The x-axis shows the percentage of cortical vertices assigned to each network. The y-axis shows the percentage of cerebellar voxels assigned to each network in the contralateral hemisphere. The black lines indicate best-fit regression lines.

In general, networks occupying larger proportions of the cerebral cortex possessed larger cerebellar representations. Across the two subjects and including both hemispheres, the relationships ranged from r = 0.41 to 0.81. The results did not reveal any consistent pattern that might suggest disproportionate representation of a single association network in the cerebellum. In particular, the two networks associated with cognitive control were represented as expected in the cerebellum, with no clear disproportionate relationship as compared to the default network A and B, the language network, or the dorsal attention network.

## Discussion

Detailed analysis of cerebellar organization reveals a macroscale organization. Within this broad macroscale organization are additional reproducible topographic features that include separation of distinct higher-order association networks linked to the default network, language, and cognitive control.

### A macroscale organization is present in the cerebellum despite individual differences

A key result of the present findings is that a conserved topographic organization of the cerebellum emerges reproducibly between individuals despite a complex topography and individual differences. Both intensively-sampled individuals analyzed here demonstrated the same broad macroscale organization. These findings support the hypothesis that, just as the cerebral cortex demonstrates a broad macroscale organization, the topography of the cerebellum is likely driven by developmental constraints – a conserved *Bauplan* – that is expressed between individuals despite differences in idiosyncratic anatomical features.

In many ways this result is expected. Global organizational features such as the double somatomotor map with its inverted orientation in the anterior lobe and upright orientation in the posterior lobe are conserved between distantly related mammalian species including cats and monkeys (Adrian 1943; Snider and Stowell 1944) and detected using direct physiological methods (Adrian 1943; Snider and Stowell 1944), task-based functional MRI (Grodd et al. 2001; Wiestler et al. 2011; Buckner et al. 2011; Diedrichsen and Zotow, 2015; Boillat, Bazin and van der Zwagg 2020), and intrinsic functional connectivity (Buckner et al. 2011; Marek et al. 2018). The present data extend these observations to suggest that the general spatial arrangements of the cerebellar association zones are conserved between individuals. Specifically, the network organization in the cerebellum follows a hierarchical macroscale gradient paralleling recent observations in the cerebral cortex (Margulies et al. 2016; Krienen and Buckner 2017; Hutenberg et al. 2017; Buckner and DiNicola 2019).

The primary cerebellar hierarchy begins in the anterior lobe, progressing from the primary somatomotor zone to first order association networks linked to sensory-motor processes (the dorsal attention and salience/ventral attention networks) and then to higher-order association networks (cognitive control and default network A and B). The hierarchy reverses and progresses through the same set of networks in opposite order, ending in the secondary somatosensory zone in VIIIa/VIIIb. This major gradient is consistent with that proposed by Guell et al. (2018b). Like the cerebral hierarchy, the cerebellar zones progress from networks involved in processing and acting on the external sensory environment to anatomically distant zones linked to networks involved in constructive aspects of cognition. Why there are major gradients in the cerebellum, and why they show an inverted topography mirrored on either side of the Crus I/II border are open questions that will likely be informed by better understanding of developmental mechanisms that specify early projection patterns.

Evidence for a third hierarchical representation situated in the most posterior extent of the cerebellum is also provided, as hypothesized based on group data (Buckner et al. 2011; Guell et al. 2018b). In both individuals, clear representations of default network A and B were present in IX, surrounded by representations of the cognitive control network anteriorly and then the dorsal attention and salience/ventral attention networks before the prominent somatomotor representations in VIIIa/VIIIb. The posterior representations of default network A and B in IX, when seeded with small regions, were each sufficient to reproduce the full distributed extents of the separate association networks (Fig. 12). The possibility of a posterior representation of cognitive networks was anticipated by the polysynaptic tracing observations of Kelly and Strick (2003) who observed projections to and from the prefrontal cortex to lobule IX/X in the monkey (see their Fig. 11). Here we reveal that these posterior zones, when analyzed within the individual, possess distinct localized representations of the multiple association networks.

### A cerebellar language network can be distinguished from adjacent association networks

One of the earliest observations suggesting the involvement of the cerebellum in cognition was that the right lateral cerebellum was active during cognitive demands of a speech production task (Petersen et al. 1989). Despite this origin, multiple published estimates of the cerebellar organization based on group functional connectivity MRI (Buckner et al. 2011) and individual functional connectivity MRI (Marek et al. 2018) do not identify a clear language network in the cerebellum. A contributing reason for this omission is likely that networks important to language are adjacent to the largest association network identified with functional connectivity (the default network) and are blurred together in many functional connectivity analyses (for discussion, see Braga et al. 2020).

Recent cerebellar mapping efforts explicitly using tasks demanding language processing yield clear responses at or near the Crus I/II border (King et al. 2019; see also Guell et al. 2018a). For example, using a verb generation task similar to that originally employed by Petersen and colleagues, King et al. (2019) demonstrated a large region of response near right Crus I/II that overlapped the response observed during a theory-of-mind task (see their Fig. 1). These prior studies make clear that networks important to language are represented in the cerebellum, but also that the location of the regions and their separation from other networks is challenging. Within the cerebrum, specialized regions involved in language are finely juxtaposed next to distinct higher-order association regions involved in cognitive control and default network A and B (Fedorenko et al. 2012; Braga et al. 2020).

Here we found clear representation of a language network in the cerebellum that is spatially separate from nearby association networks. In both individuals, the language network abutted the major Crus I/II zones linked to default network B and cognitive control network A, echoing their close proximity in the cerebral cortex (Braga et al. 2020). In one individual the network was strongly right lateralized and in the other more bilateral, reflecting variation in lateralization that is well known in the cerebrum (and provisionally established for the cerebellum; Wang, Buckner and Liu 2013). These findings suggest the presence of multiple discrete cerebellar zones that are components of a network specialized for language. The cerebellar language network regions fall within a larger zone of multiple, specialized association networks, as discussed in the next section.

### The Crus I/II apex possesses juxtaposed zones that participate in specialized networks

Beyond the broad gradient evident in both individuals, at the Crus I/II border, which forms the apex of the cerebellum, multiple association networks were juxtaposed next to one another with idiosyncratic spatial patterns between individuals. The flatmap visualization in Fig. 10 displays this feature. Virtually all networks have multiple representations that fall along the medial to lateral extent of the cerebellum, which are interdigitated with distinct association networks. Detailed analyses of the focal zones in Figs. 12-14 support that most of these local cerebellar zones are specific for a single cerebral network rather than a technical artifact of a winner-take-all parcellation. For example, the cerebellar zones that are associated with the language network possess multiple representations at the edges of the Crus I / II association cluster. Each of these representations, when seeded in the cerebellum, yields the distributed cerebral association network linked to language function (Fig. 13) and can be dissociated from nearby zones (Fig. 14). As another example, the local cerebellar zones that are linked to default network A and B (Braga and Buckner 2017; Braga et al 2019; DiNicola et al. 2020) show multiple dissociated representations in the cerebellum, including separate medial and lateral components (Fig. 12). The organization of these multiple and interdigitated representations of networks within the Crus I/II association region was not revealed in earlier group-averaged analyses of cerebellar topography and do not appear to reflect a simple linear gradient.

The complex organization within individuals raises several issues to consider for future studies. First, description of the detailed organization of the cerebellum requires high resolution, almost certainly better than that achieved here or in the field to date. Features of functional organization are missed when group-averaged or lower-resolution estimates are obtained. Second, there are translational implications of the present results. The Crus I/II apex is the closest portion of the cerebellum to the scalp (∼15mm), approximately half the distance to the estimated somatomotor representations in lobule V (∼30-35mm). Given that the cerebellum has emerged as a neuromodulatory target for neuropsychiatry (e.g., Brady et al. 2019) and transcranial magnetic stimulation (TMS) can alter cognitive domains (Desmond, Chen and Shieh 2005; Esterman et al. 2017; Sheu, Liang and Desmond 2019), detailed modeling of stimulation effects on distinct networks at and around the Crus I/II apex will become increasingly relevant. The juxtaposition of network regions and individual variation will be important considerations.

Finally, the complex interdigitation of association networks – as an anatomical feature – is itself a target for understanding. Similar to the cerebral cortex, the cerebellum possesses a broad macroscale organization (see also Buckner et al. 2011; Guell et al. 2018a). Within that organization is a set of fractionated zones linked to distinct cerebral networks, supporting specialized cognitive functions. At the Crus I/II apex there are zones linked to cerebral networks involved in language, remembering, and aspects of social cognition. Zones of the cerebellum supporting these advanced forms of cognition and the boundaries between them seem unlikely to be fully specified by innate molecular gradients. Rather, paralleling what has been hypothesized for cerebral organization (e.g., Buckner and DiNicola 2019; DiNicola et al. 2020), cerebellar development may be influenced by protracted activity-dependent sculpting that differentiates these interdigitated zones within an earlier-specified broad gradient. A hierarchical developmental sequence might explain the broad macroscale organization that is observed across individuals (and even species) as well as the interdigitated and varied organization that exists within regions including the Crus I/II apex (see also Marek et al. 2018).

### The cerebellar vermis represents the central field of primary visual cortex

Visual topography at or near the vermis has not widely been discussed in prior studies of human cerebellar organization, including in our own work (e.g., Buckner et al. 2011). However, motivated by the present work, we revisited our earlier group-averaged cerebellar parcellations and did find that the vermis was coupled to a visual network representing peripheral visual eccentricities (Fig. 17). More recently, in a thorough and detailed analysis of the vermis using visually-evoked task data from the HCP, van Es and colleagues (2019) reported strong evidence that the vermis maps to topographically organized visual space. The present results provide convergent evidence that the vermis is coupled to a cerebral visual network and further reveals that the mapping (1) is linked to primary visual cortex and (2) possesses representation of the central visual field.

An additional feature of the vermis mapping is that it can be distinguished from a nearby but separate cerebellar representation of the dorsal attention network (Fig. 16). The two sets of regions within the cerebellum are coupled to non-overlapping networks in the cerebral cortex. The vermis is coupled to early retinotopic cortex that includes central V1; distributed zones that include prominent representations in lateral VI and VIIb/VIII are coupled to the dorsal attention network, consistent with the observations of Brissenden and colleagues (2018).

Boillat and colleagues (2020) recently generated human task-based somatomotor maps in the cerebellum at high resolution within the individual through instructed movement of body parts, including the eyes. In their eye movement task, a dot jumped back and forth across the screen and the subjects followed it. Their analysis revealed an eye-selective response within the vermis (see also King et al. 2019). As van Es et al. (2019) note in their detailed analysis of the vermis, maps based on retinotopic stimulation while subjects are fixating cannot unambiguously distinguish among retinotopic (eye-centered), head-centered, and world-centered spatial mapping. It is thus of particular interest that the present results identify the vermis of the cerebellum through its functional coupling to early retinotopic visual cortex, in the cerebellar region that has also been identified in studies of eye movement.

Work in non-human primates has long suggested that the cerebellar vermis participates in spatially-mapped oculomotor function including adaptive response to saccadic errors (Optican and Robinson 1980; Sparks and Mays 1990; Herzfeld et al. 2015; Soetedjo et al. 2019). For this reason, the region has often been labeled the oculomotor vermis or OMV (e.g., Yamada and Noda 1987; van Es et al. 2019). Focus of OMV’s role in saccade error correction has most often highlighted signals originating from the superior colliculus and on aspects of motor-control related to saccadic adjustment. The present results suggest OMV, directly or indirectly, represents information coupled to primary visual cortex (Figs. 16-19). The finding that OMV is coupled to primary visual cortex, while incompletely understood at this time, may be relevant to understanding how motor predictions are integrated with their perceptual outcomes.

### Limitations and caveats

A key limitation of the present results is that they are based on correlations between cerebral and cerebellar signals. Future work will be required to verify the functional-anatomical details through differential task-induced responses of the cerebellar subzones proposed to be distinct via the present findings. In this light, the detailed maps of the cerebellum generated here within individual subjects should be considered hypotheses, much like our earlier work on group averaged data (Buckner et al. 2011) and those from other efforts (Stoodley and Schmahmann 2009; Guell et al. 2018a; Marak et al. 2018; King et al. 2019). The present work adds to this progressing endeavor by describing new details around the Crus I/II border where multiple, previously undistinguished zones, are demonstrated to couple to distinct interdigitated cerebral networks.

A further limitation of the present work, that also applies to prior functional neuroimaging work on the human cerebellum, pertains to resolution. The folia of the cerebellum are organized like a fine-folded accordion that, if unraveled, are about 1m long and 10cm wide (Sereno et al. 2020; see also Sultan and Braitenberg 1993; van Essen 2002; Diedrichsen and Zotow 2015). Small errors in localization in the volume can translate into differences of centimeters on the modeled cerebellar surface. Our functional data collected here at 2.4mm resolution and smoothed during processing falls far short of what would be required to resolve the cerebellar cortex (Sereno et al. 2020 used <0.2mm voxels to reconstruct the cerebellar surface). For the foreseeable future, functional analyses within individuals will be based on probabilistic relationships to the details of surface geometry. The present analyses likely miss (or distort) topographic features of functional anatomy (see Diedrichsen and Zotow 2015; Sereno et al. 2020 for discussion).

One technical point to mention for future experimentalists is that the present results were not our first attempt to map within-individual organization of the cerebellum. Beginning shortly after our initial efforts to map the cerebellum in the group (Buckner et al. 2011), we attempted several times to map the detailed organization, especially the hypothesized third map near the posterior lobe (the 7T reported in Braga et al. 2019 was originally analyzed for such a purpose). In each instance, while aspects of cerebrocerebellar topography could be mapped using functional connectivity (e.g., the representation of the default network in Crus I / II), other aspects needed for validation failed (e.g., the detection and reversal of the body maps as illustrated here in Fig. 3). The clear patterns presented in this paper only emerged when we collected extremely large quantities of data within the same individual, thereby boosting signal-to-noise significantly. We do not yet know the boundary conditions of what amount of data are required to explore within-individual anatomy at high resolution and how alternative analytical approaches might improve the situation, but we have been struck by the need to acquire far more data than was necessary for our parallel analyses of the cerebrum.

### Conclusions

The human cerebellum possesses a topographic representation of multiple higher-order association networks. Separate cerebellar zones can be shown to couple selectively to distinct networks that are finely interdigitated in the cerebral cortex, including networks hypothesized to support advanced functions related to remembering, making social inferences, and language.

## Acknowledgements

We thank R. Mair, A. Youssoufian, H. Becker and E. Phlegar for assisting in data acquisition, T. O’Keefe for preprocessing optimization, and N Saadon-Grosman for insights into the organization of the body maps. We thank the Harvard Center for Brain Science neuroimaging core and FAS Division of Research Computing. The multi-band EPI sequence was generously provided by the Center for Magnetic Resonance Research (CMRR) at the University of Minnesota.

